# Cognitive boundary signals in the human medial temporal lobe shape episodic memory representation

**DOI:** 10.1101/2021.01.16.426538

**Authors:** Jie Zheng, Andrea Gómez Palacio Schjetnan, Mar Yebra, Clayton Mosher, Suneil Kalia, Taufik A. Valiante, Adam N. Mamelak, Gabriel Kreiman, Ueli Rutishauser

## Abstract

While experience unfolds continuously, memories are organized as a set of discrete events that bind together the “where”, “when”, and “what” of episodic memory. This segmentation of continuous experience is thought to be facilitated by the detection of salient environmental or cognitive events. However, the underlying neural mechanisms and how such segmentation shapes episodic memory representations remain unclear. We recorded from single neurons in the human medial temporal lobe while subjects watched videos with different types of embedded boundaries and were subsequently evaluated for memories of the video contents. Here we show neurons that signal the presence of cognitive boundaries between subevents from the same episode and neurons that detect the abstract separation between different episodes. The firing rate and spike timing of these boundary-responsive neurons were predictive of later memory retrieval accuracy. At the population level, abrupt neural state changes following boundaries predicted enhanced memory strength but impaired order memory, capturing the behavioral tradeoff subjects exhibited when recalling episodic content versus temporal order. Successful retrieval was associated with reinstatement of the neural state present following boundaries, indicating that boundaries structure memory search. These findings reveal a neuronal substrate for detecting cognitive boundaries and show that cognitive boundary signals facilitate the mnemonic organization of continuous experience as a set of discrete episodic events.

## Introduction

Our lives unfold over time, weaving rich, dynamic, and multisensory information into a continuous sequence of experiences. However, our memories are not continuous. Rather, what we remember are discrete episodes (“events”)^1^, which serve as anchors to bind together the myriad different aspects (where, when, what) of a given autobiographical memory^2^ much like objects do in perception^3^. For instance, the mnemonic representation of a movie consists mainly of a set of salient moments, disregarding large amounts of other information and irrespective of their temporal order in the original movie^4^. A fundamental unresolved question in human memory is, therefore: what marks the beginning and the end of an episode?

The transformation from ongoing experience to distinct events is thought to rely on the identification of boundaries that separate two events^1,5–9^. Neuroimaging studies in humans indicate that neural activity in the medial temporal lobe (MTL), in particular in the hippocampus and parahippocampal gyrus, changes around the occurrence of cognitive boundaries and the extent of such changes is related to later memory performance^10,11^. However, due to the low temporal and spatial precision, it remains unclear how exactly these responses are related to abstract cognitive boundaries, when these responses occur, and what the mechanisms of their generation are. In rodents, much has been learned from studying the neural responses to spatial boundaries and the ways by which these responses shape mnemonic representations. For example, neurons in the rodent hippocampus elevate their firing rates in the vicinity of investigator-imposed spatial boundaries^12^, with the place fields of hippocampal neurons shaped by physical boundaries like turns^13^ and walls^14,15^. In accordance with the boundaries of subenvironments^12^, hippocampal place fields remap^16,17^ in response to context shifts (e.g., enter a new compartment) and are reinstated^13,18^ when placed back to a familiar context. Additionally, rodent MTL neuron ensembles encode event-specific representations irrespective of an animal’s spatial location^19^, presumably representing cognitive boundaries between distinct events. Together, boundaries shape mnemonic representations of both spatial environments and the events that occur along the way, and structure the neural basis (i.e., place fields and event-specific representations) of cognitive maps. However, no such understanding exists yet for the non-spatial episodic memories that define us as individual human beings^2,20^.

We investigated the neuronal mechanisms underlying the identification of event boundaries in humans under relatively realistic continuous experience. We recorded single neuron activity from 20 patients with drug-resistant epilepsy with implanted depth electrodes^21^ while testing their memory for the content of video clips with two different kinds of embedded cognitive boundaries: soft and hard boundaries. Soft boundaries are episodic transitions between related events within the same movie, while hard boundaries are episodic transitions between unrelated events from two unrelated movies. Behaviorally, we found that both soft and hard boundaries enhanced recognition of video clip content that followed a boundary, whereas hard boundaries impaired memory of the temporal order between events. This tradeoff is compatible with the segmentation of experience into distinct episodes. Neuronally, we characterized the properties of neurons in the medial temporal lobe that signaled the presence of boundaries. The activity of these boundary responsive neurons predicted memory strength as assessed by scene recognition and temporal order discrimination accuracy.

## Results

We studied how boundaries influence the formation and retrieval of memories of brief video clips. Twenty patients with drug-resistant epilepsy implanted with hybrid depth electrodes for localizing their seizure foci performed the task while we recorded the activity of single neurons (Fig. 1e; Extended Data Tables 2 and 3 show electrode locations and patient demographics). The task consisted of three parts: encoding, scene recognition, and time discrimination. During encoding (Fig. 1a), subjects watched 90 different and novel video clips containing either no boundaries (NB, one continuous movie shot), soft boundaries (SB, cuts to a new scene within the same movie), or hard boundaries (HB, cuts to a new scene from a different movie; See Fig. 1b and Extended Data Fig. 1 for examples of the different types of boundaries). Differentiating between SB and HB required cognitive understanding of movie content because visual features (Extended Data Table 1) did not differentiate between these two types of boundaries. To ensure subjects’ engagement during encoding, a memory question (e.g., is anyone in the clip wearing glasses?) appeared every four to eight clips. Subjects answered 89 ± 5% of these questions accurately.

**Fig. 1:**
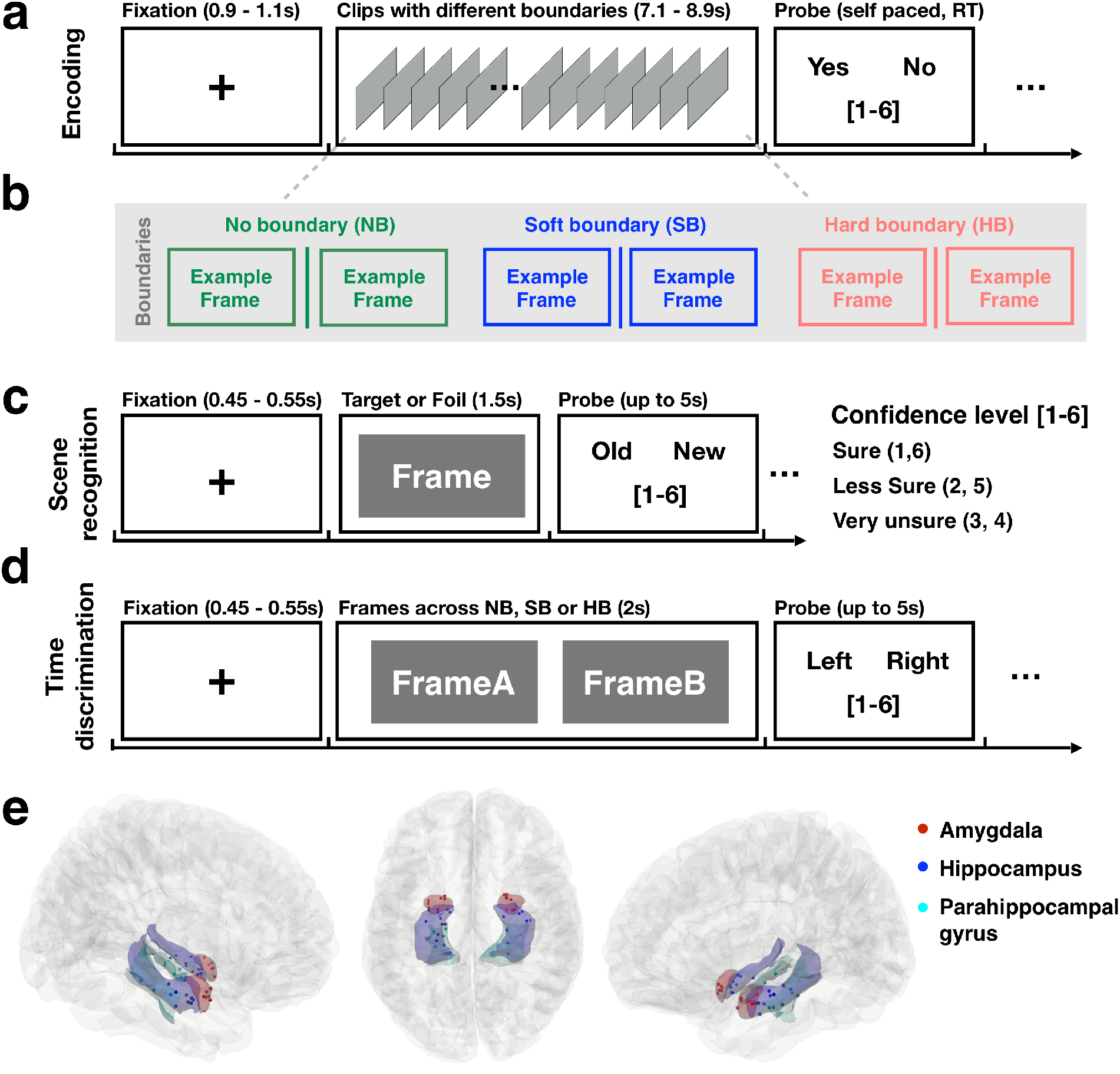
Experiment and recording locations. **a.** Encoding task (example trial). Subjects watched 90 video clips (~ 8 seconds each, no audio) with either no boundary (NB, continuous movie shot), a soft boundary (SB, cut to a new scene within the same movie, 1 to 3 SB per clip), or a hard boundary (HB, cut to a different movie, 1 HB per clip). Every 4-8 clips, subjects were prompted to answer a true/false question related to the clip content together with a confidence rating (see Methods). **b**. Example boundaries (more examples in **Extended Data Fig.1**, visual features of boundaries in **Extended Data Table 1**). Note that owing to copyright issues, all original images have been removed but are available upon reasonable request. **c**. Recognition memory task. Subjects indicated whether a static image was new or old (seen in the video clips), together with a confidence judgment. **d**. Time discrimination task. Subjects indicated which of two frames they saw first during the movie clips together with a confidence rating. **e.** Recording locations of 39 electrodes (see MNI coordinates in **Extended Data Table 2**) across all subjects (subject information in **Extended Table 3**) in the amygdala (red), hippocampus (blue), or parahippocampal gyrus (cyan), rendered on the Colin27 template brain^14^. Each dot represents the location of a microwire bundle.

We subsequently evaluated what subjects remembered about the video clips with two memory tests: scene recognition (Fig. 1c) and time discrimination (Fig. 1d). During the scene recognition test, subjects were presented with individual static frames. These frames were chosen with equal probability from either the previously presented NB, SB, and HB video clips (“targets”) or from other video clips that were not shown to the subjects (“foils”). Subjects made an “old” or “new” decision together with a confidence rating (sure, less sure and very unsure) for each. During the time discrimination test, subjects were shown two old frames from the video clips side-by-side (Fig. 1d) and had to indicate which frame was seen earlier in time together with a confidence rating.

### Boundaries strengthen recognition memory but disrupt temporal order memory

In the time discrimination task, subjects correctly identified which frame was shown first in 73 ± 7% and 73 ± 8% of trials when the two frames were separated by no boundary or a soft boundary in the video clips, respectively (Fig. 2a, green and blue; both above chance of 50%; NB: p < 0.001; SB: p< 0.001). In contrast, subjects performed significantly worse when discriminating between frames that were separated by a hard boundary (Fig. 2a, red, HB: 53% ± 5% vs 73% for NB and SB; *F* (2, 57) = 51.33, *p* = 1.78×10^−13^; p = 0.02 against chance level). Subjects also showed longer reaction times (Fig 2b; HB: 2.10 ± 0.37 seconds; NB: 1.62 ± 0.28 seconds; SB: 1.59 ± 0.34 seconds; *F* (2, 57) = 14.25 *p* = 9.6×10^−6^) and lower confidence ratings when discriminating between two frames earlier separated by a hard boundary (Fig 2c; HB: 1.95 ± 0.45; NB: 2.52 ± 0.29; SB: 2.59 ± 0.23; *F* (2, 57) = 20.41, *p* = 2.07×10^−7^). This effect on RT and confidence was not driven by accuracy differences as it was observable for both correct and incorrect trials independently (see Extended Data Fig. 3).

**Fig. 2:**
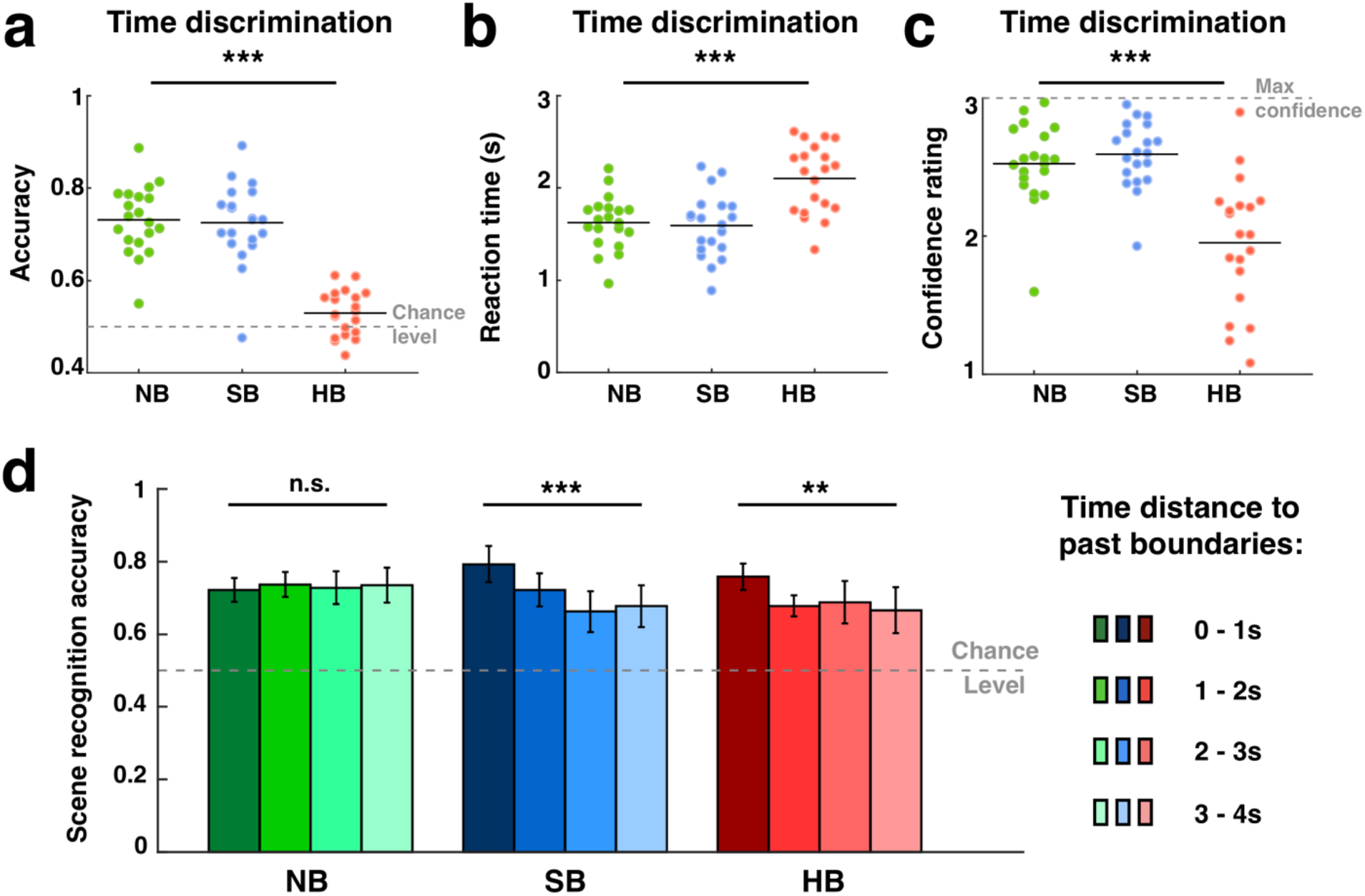
Behavior. Hard boundaries impaired time discrimination while soft and hard boundaries improved recognition memory for frames close to them. **(a-c)** Accuracy (**a**), reaction time (**b**), and confidence level (**c**) across all the trials for NB (green), SB (blue), and HB (red) during the time discrimination task (see also **Extended Data Fig. 3–4**). The horizontal dashed lines in (**a**) show chance levels (0.5) and in (**c**) they show the maximum possible confidence rating (max = 3). Behavior data for the scene recognition task is shown in **Extended Data Fig. 2**. **d.** Scene recognition accuracy as a function of boundary type and time elapsed between the target frame and its nearest past boundary (distance effect for time discrimination accuracy and future boundaries shown in **Extended Data Fig.5**). In the NB clips, we measured time with respect to the middle of the clip. Error bars indicate standard deviation across n= 20 sessions. \*\**P* < 0.01, \*\*\**P* < 0.001, one-way ANOVA, degrees of freedom = (2, 57) for (**a-c**) and (3, 76) for (**d**).

Across all trials, the ability to recognize a frame as ‘old’ did not differ significantly between the type of boundary that preceded the frame (Extended Data Fig. 2a; NB: 76% ± 10%; SB: 75% ± 9%; HB: 75% ± 8%, *F* (2, 57) = 0.07, *p* = 0.94). The reaction times and confidence ratings during the scene recognition task were also similar across the different types of boundaries (reaction time: Extended Data Fig. 2b; NB = 1.47 ± 0.18 seconds, SB = 1.43 ± 0.16 seconds, HB = 1.49 ± 0.15 seconds*, F* (2, 57) = 0.28 *p* = 0.76; confidence rating: Extended Data Fig. 2c; NB = 2.60 ± 0.18, SB = 2.60 ± 0.20, HB = 2.52 ± 0.28, *F* (2, 57) = 0.54, *p* = 0.56). Therefore, the impaired time discrimination ability across HB transitions was not due to differences in memory strength as measured in recognition accuracy. Even though the overall accuracy was similar among NB, SB, and HB conditions, the recognition accuracy of target frames decreased as a function of how far away in time the target frame was relative to the immediately preceding boundary. Target frames presented within 1s after a SB (accuracy = 79% ± 6%) were remembered better than those further away from the boundary (Fig. 1d, blue; 1-2s: 72% ± 6%; 2-3s: 66% ± 9%; 3-4s: 68% ± 9%; *F* (3, 76) = 10.55, *p* = 7.02×10^−6^). A similar effect was also present following HBs: target frames presented within 1s after a HB (accuracy = 76% ± 5%) were significantly better remembered than those farther away from the boundary (Fig. 1d, red; 1-2s: 68% ± 4%; 2-3s: 69% ± 9%; 3-4s: 67% ± 10%; *F* (3, 76) = 5.47, *p* = 0.002). In contrast, recognition accuracy did not differ significantly as a function of time in the no boundary condition (Fig. 1d, green; 0-1s: 72% ± 5%; 1-2s: accuracy = 74% ± 5%; 2-3s: accuracy = 73% ± 6%; 3-4s: accuracy = 74% ± 7%; *F* (3, 76) = 0.32, *p* = 0.81). The distance effects described were unidirectional because the distance to future boundaries did not influence memory performance (Extended Data Fig. 5ab). No distance effect was present for time discrimination accuracy (Extended Data Fig. 5cd).

Together, behavioral analysis revealed that frames that closely followed a soft or hard boundary were more likely to be later recognized. Memory for the temporal order of frames, on the other hand, was disrupted by the presence of hard boundaries. Hard boundaries were thus beneficial for recognition memory but detrimental for order memory – thereby revealing a fundamental trade-off.

### Neurons in the medial temporal lobe demarcate episodic transitions

We next investigated the neuronal responses to boundaries and their relationship to memory by recording from neurons in the medial temporal lobe (MTL; we recorded in the hippocampus, amygdala and parahippocampal gyrus) as well as other brain areas (Extended Data Tables 2 and 4). Across all areas, we recorded the activity of 985 neurons (Extended Data Fig. 6 shows spike sorting quality) from 19 subjects (1 of the 20 subjects yielded no usable recordings, see Extended Data Table 3). We first tested whether neurons changed their activity following the occurrence of boundaries by selecting for neurons that increased their response in a 1s long window following boundaries (i.e., post-boundary responses) relative to baseline (1s prior to boundary) in SB and HB trials, but not in NB trials (see Methods). Fig. 3a-b shows two of the selected neurons recorded from the parahippocampal gyrus and hippocampus, respectively. These neurons showed a transient increase in firing rates within approximately 300 milliseconds after both soft (blue) and hard (red) boundaries. We refer to this type of neuron as a boundary cell. Forty-two out of the recorded 580 MTL neurons (7.24%; chance = 2.11%) were classified as boundary cells (Fig. 3c), with the largest proportions located in the parahippocampal gyrus (n= 18/68, 26.47%), followed by amygdala (n= 12/169, 7.10%) and hippocampus (n= 12/343, 3.50%; see Extended Data Table 4 for statistics).

**Fig. 3:**
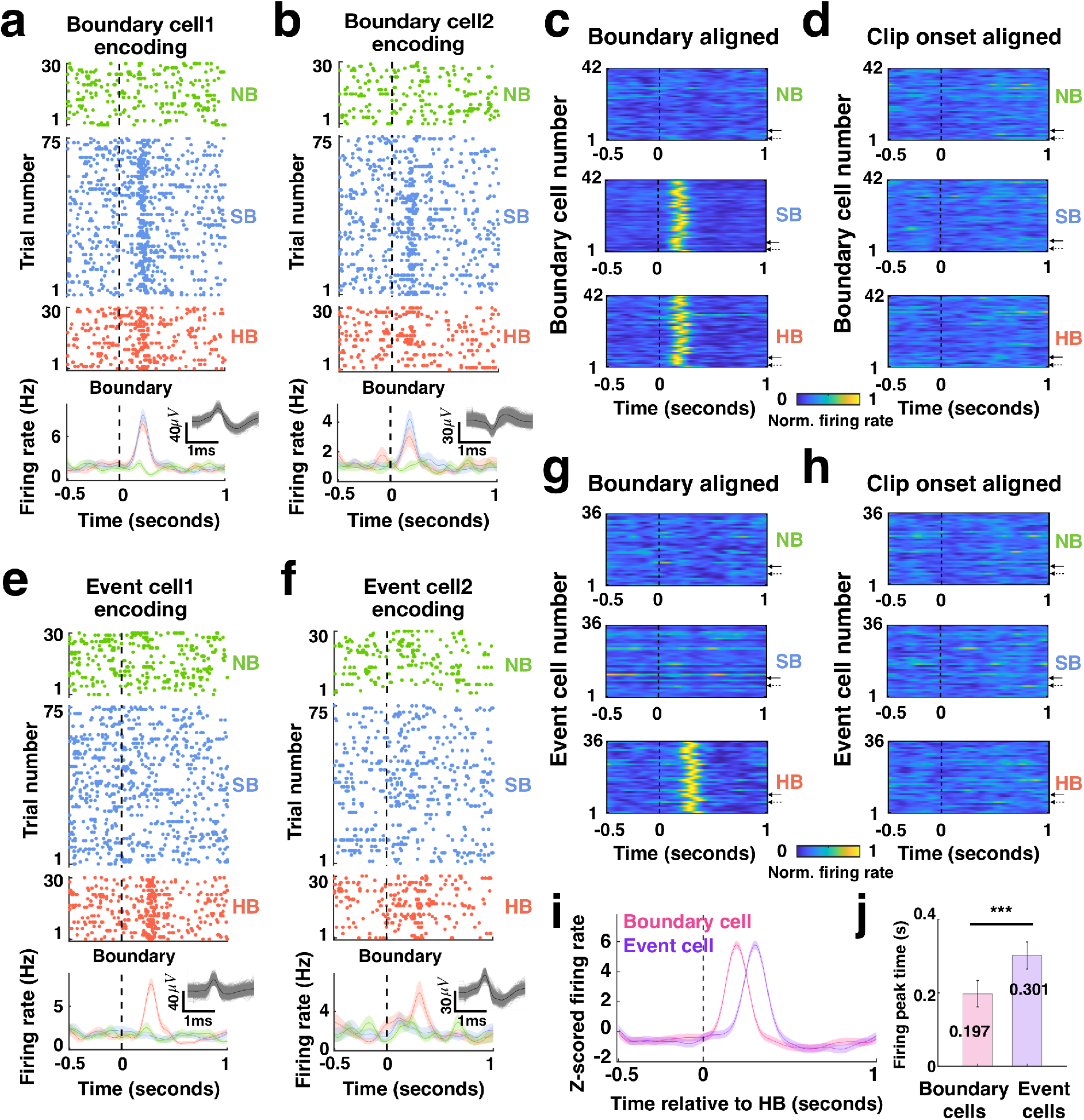
Boundary cells and event cells demarcate different types of episodic transitions. **a**-**b.** Responses during the encoding stage from two example boundary cells located in the parahippocampal gyrus and hippocampus, respectively (spike sorting quality of all detected cells shown in **Extended Data Fig. 6**). Boundary cells responded to both SB (blue) and HB (red) transitions. Responses are aligned to the middle point of the clip (NB, green) or to the boundary (SB, HB). Top: raster plots. Bottom: Post-stimulus time histogram (bin size = 200 ms, step size = 2ms, shaded areas represented ± s.e.m. across trials). Insets: all spike extracellular waveforms (gray) and mean (black). **c-d**. Firing rates of all 42 boundary cells (solid and dashed arrows denote the examples in **a** and **b**, respectively) during the encoding stage aligned to the clip onsets (**d**) or boundaries (**c**), averaged over trials within each boundary type and normalized to each neuron’s maximum firing rate from the entire task recording (see color scale on bottom). Boundary cells responded to boundaries (SB and HB) (**c**) but not to clip onsets (**d**) or clip offsets (**Extended Data Fig.7a-c)**. **e**-**f.** Responses during the encoding stage from two example event cells located in the hippocampus and amygdala, respectively. Event cells responded to HB (red) but not SB (blue) transitions. **g-h**. Firing rates of all 36 event cells (solid and dashed arrows denote the examples in **e** and **f**, respectively) during the encoding stage, using the same format as **c-d**. Event cells responded to HB (**g**) but did not respond to clip onsets (**h**) or clip offsets (**Extended Data Fig.7d-f)**. **i.** Latency analysis. Firing rate during HB transitions (to which both boundary cells and event cells responded) reached peak response earlier for boundary cells (pink) compared to event cells (purple). Shown is average z-score firing rate normalized using the average and standard deviation of the firing rates and aligned to HB (bin size = 200 ms, step size = 2ms, shaded areas represented ± s.e.m. across all boundary cells or event cells). **j**. Peak times of average firing rate traces of all boundary cells (pink) and all event cells (purple). Error bars indicate standard deviation across n = 42 or 36 cells, respectively. \*\*\**P* < 0.001, one-way ANOVA, degrees of freedom = (1, 76). The spatial distribution of boundary cells and event cells is shown in **Extended Data Table 4**. See **Extended Data Fig. 8–9** for responses of boundary cells and event cells during the scene recognition and time discrimination stages.

Was the response of boundary cells a result of the abrupt changes in pixel-level content from the frame before to the frame after the boundary? To answer this question, we considered the responses of the cells during two other abrupt changes of visual input: video clip onset (Fig. 3d) and offset (Extended Data Fig. 7). Boundary cells did not respond significantly to video clip onset or offset (p > 0.05; permutation t-test, see Methods), showing that the boundary-related response of boundary cells is likely related to higher level cognitive discontinuities rather than pure visual changes (a question we return to below).

We also found a second group of neurons that transiently increased their firing rate only following hard, but not soft boundaries or no boundaries (see Methods). Two examples of such cells, located in the hippocampus and amygdala, are shown in Fig. 3e-f. We refer to this type of response as an event cell. Thirty-six out of the recorded 580 MTL neurons (6.20%; chance = 2.26%) were classified as event cells (Fig. 3g), with the largest proportions located in the hippocampus (n= 27/343, 7.87%), followed by amygdala (n= 7/169, 4.27%), and parahippocampal gyrus (n= 2/68, 2.94%). Similar to boundary cells, event cells did not significantly change their firing rates (p > 0.05, permutation t-test; see Methods) following video clip onset (Fig. 3h) or offset (Extended Data Fig. 7c, f).

The types of transitions that we refer to as soft and hard boundaries differ in terms of their high-level conceptual narrative, which is interrupted in HBs but not in SBs. Is it possible to determine from visual features alone whether a boundary was of the soft or hard kind? To assess this question, we computed the differences between pre- and post-boundary frames in terms of pixel-level characteristics (i.e., luminance, contrast, complexity, entropy, color distribution), high-level visual features (i.e., objects), and perceptual similarity ratings. These analyses revealed that SB and HB transitions did not differ significantly from each other in any of the attributes we tested (see Extended Data Table 1). Therefore, the differential activation of event cells to HBs but not SBs is likely a result of detection of the disruption in the conceptual narrative, that is, a transition between two different episodes.

While both boundary and event cells responded to HB type transitions, a comparison of their response dynamics indicated that boundary cells responded to hard boundaries approximately 100 milliseconds earlier than event cells (Fig. 3i). This effect was also visible when comparing the time at which their peak response was reached: boundary cells showed a peak at 197 ± 49 milliseconds, whereas event cells showed a peak at 301 ± 55 milliseconds (Fig. 3j; *F* (1, 76) = 17.71, *p* = 7×10^−5^).

We also selected for boundary and event cells in brain areas other than the medial temporal lobe, such as the medial frontal cortex, insula, and orbitofrontal cortex (OFC). We found 8/405 (1.96%) boundary cells and 9/405 (2.22%) event cells among the non-MTL group (see Extended Data Table 4), with only event cells in the OFC exceeding the proportions expected by chance. These results show the specificity of boundary responsive neurons to the MTL; we restrict the following analyses to the MTL neurons.

### The responses of boundary and event cells predict later memory strength

We next asked whether the responses of boundary and event cells during video watching (encoding) correlated with later measures of memory for the content of the videos. We examined whether the strength of responses of boundary cells or event cells to boundaries varied as a function of whether the familiarity or temporal order of a stimulus was later remembered or forgotten. Fig. 4a show an example boundary cell located in the hippocampus whose response during encoding differed between video clips from which frames were later correctly remembered as familiar (Fig. 4a_1_) vs. incorrectly identified as novel (forgotten, Fig. 4a_2_): the responses to boundaries that preceded later remembered frames was significantly stronger. This effect was present, on average, in the group of previously selected boundary cells (n = 42) for frames proceeded by both SBs and HBs, but not by NBs (Fig. 4c; SB: *F* (1, 82) = 82.93, *p* = 4.41×10^−14^; HB: *F* (1, 82) = 156.9, *p* = 9.81×10^−21^; NB: *F* (1, 82) = 1.18, *p* = 0.28). This effect was specific to scene recognition and boundary cells. First, the firing rate of boundary cells did not significantly predict performance in the time discrimination task (Extended Data Fig. 10a, c; NB: *F* (1, 82) = 1.25, *p* = 0.27; SB: *F* (1, 82) = 1.35, *p* = 0.25; HB: *F* (1, 82) = 1.14, *p* = 0.29). Second, the firing rate of event cells (n = 36) during encoding was not predictive of recognition memory (Extended Data Fig. 11a, c; NB: *F* (1, 70) = 1.12, *p* = 0.29; SB: *F* (1, 70) = 1.63, *p* = 0.21; HB: *F* (1, 70) = 0.79, *p* = 0.38). Third, the firing rate of event cells did not predict performance in the time discrimination task (Fig. 4e shows an example and Fig. 4g shows the population summary; NB: *F* (1, 70) = 0.35, *p* = 0.56; SB: *F* (1, 70) = 0.22, *p* = 0.64; HB: *F* (1, 70) = 1.6, *p* = 0.21).

**Fig. 4:**
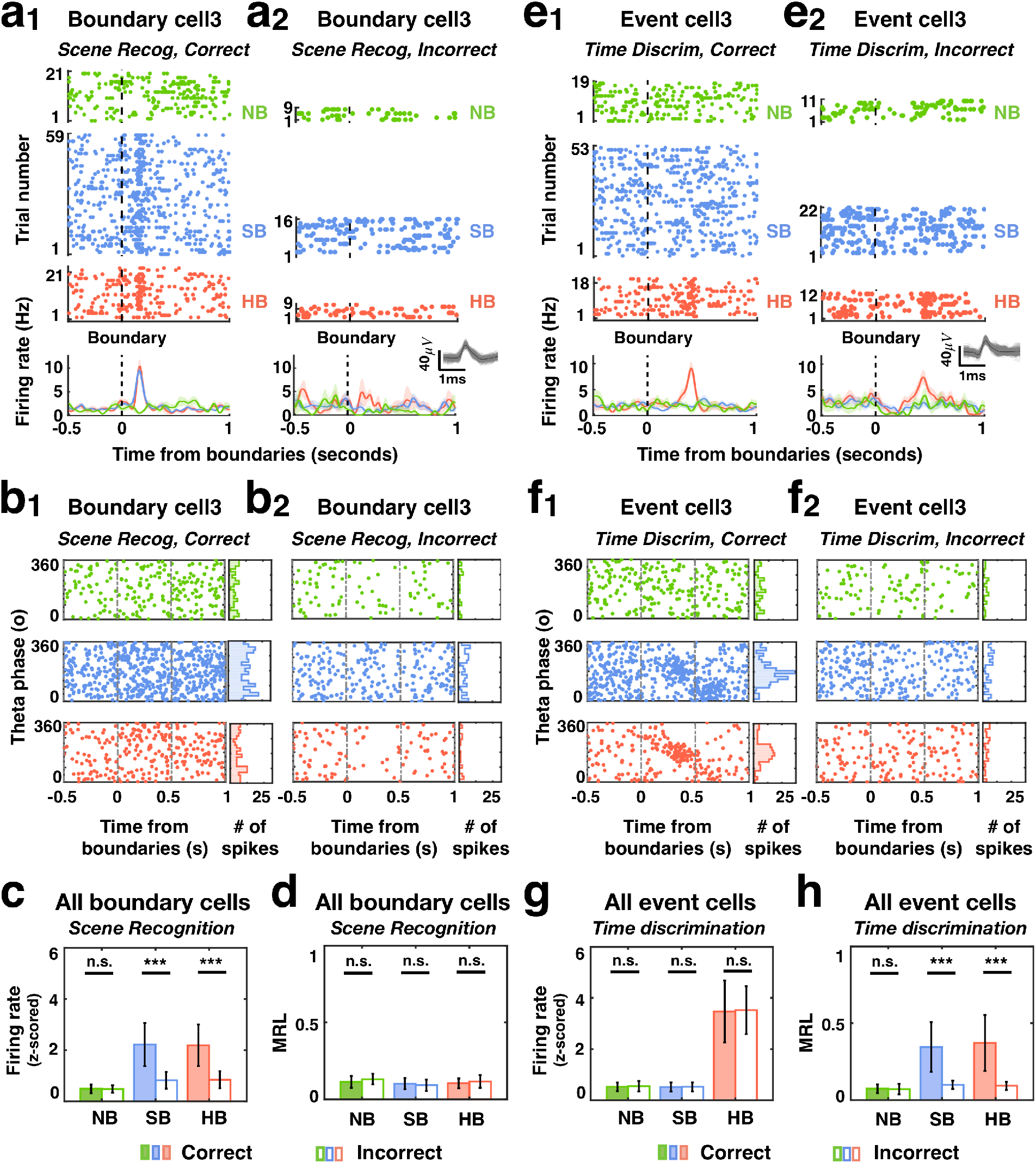
Responses of boundary cells and event cells during encoding correlate with later retrieval success. **a-d.** Response of boundary cells during encoding grouped by subjects’ subsequent memory performance in the scene recognition task. **a_1_-a_2_.** Example boundary cell recorded in the hippocampus. During encoding, this cell responded more strongly to SB and HB transitions than NB if the frame following the boundary in that trial was correctly identified during the scene recognition task (**a_1_**) compared to incorrect trials (**a_2_**). Format as in Fig. 3. **b_1_-b_2_**. Left: timing of spikes from the same boundary cell shown in **a** relative to theta phase calculated from the local field potentials, for clips of which frames were later remembered (**b_1_**) or forgotten (**b_2_**). Right: phase distribution of spike times in the 1s period following the middle of the clip (NB) or boundary (SB, HB) for clips from which frames were remembered (**b_1_**) and forgotten (**b_2_**). **c-d.** Population summary for all 42 boundary cells. **c.** Z-scored firing rate (0-1s after boundaries during encoding) differed significantly between boundaries after which frames were subsequently remembered (color filled) vs. forgotten (empty) for both SB and HB. **d**. Mean resultant length (MRL) of spike times (i.e., sum of vectors with vector lengths equal to 1 and vector angles equal to the spike timings relative to theta phases 0-1s after boundaries during encoding, divided by total number of vectors; value range [0 1]: 0 = uniform distribution, i.e., neurons fire at random theta phases; 1 = unimodal distribution, i.e., neurons fire at the same theta phase) across all boundary cells for each boundary type did not differ significantly between correct (color filled) and incorrect (empty) clips. **e-h.** Response of event cells during encoding grouped by subjects’ subsequent memory performance in the time discrimination task. **e_1_-e_2_.** Example event cell recorded in the hippocampus that responded to HB transition regardless of whether the temporal order of the clip was later correctly (**e_1_**) or incorrectly (**e_2_**) recalled in the time discrimination task. Format same as in **a** but clips were grouped based on memory outcomes in the time discrimination task. **f_1_-f_2_.** The spike timing of the same event cell shown in **e_1_-e_2_** relative to theta phase plotted for correct (**f_1_**) and incorrect (**f_2_**) trials. Format same as in **b** but clips were grouped based on memory outcomes in the time discrimination task. **g-h.** Population summary for all 36 event cells. **g.** Z-scored firing rate (0-1s after boundaries during encoding) did not differ significantly between later correctly (color filled) or incorrectly (empty) remembered temporal orders for all three boundary types. **h**. MRL of spike times (relative to theta phases, 0-1s after boundaries during encoding) was significantly larger after SB and HB transitions if the temporal order of the clip was correctly recalled (color filled) compared to the incorrect ones (empty). Error bars indicate standard deviation across n= 42 cells for **c**,**d** and n =36 cells for **g**,**h**. \*\*\**P* < 0.001, one-way ANOVA, degrees of freedom = (1, 82) for **c**,**d** and degrees of freedom = (1, 70) for **g**,**h**.

Given the importance of theta-frequency band spike field coherence in plasticity^22^, we next considered the timing of spikes with respect to theta oscillations in the local field potential (4-8Hz, measured on the microwire where the neuron was recorded from, see Methods). We determined the theta phase of each spike within a 1s window following boundaries and compared the resulting phase distributions among NB, SB and HB. This analysis revealed that event cells tended to fire at a given phase of theta following both HB and SB boundaries for clips whose temporal order was later remembered correctly (Fig. 4f shows an example). To summarize this effect across the population, we computed the mean resultant length (MRL) of all phases for all spikes fired by a given cell (see Methods). The mean MRL across all event cells (n = 36) was significantly larger following both SB and HB but not NB if temporal order was later correctly remembered (Fig. 4h; SB: *F* (1, 70) = 81.55, *p* = 2.32×10^−13^; HB: *F* (1, 70) = 60.79, *p* = 4.32×10^−11^; NB: *F* (1, 70) = 1.53, *p* = 0.22). This effect was specific to event cells and temporal order memory. First, the strength of phase-locking of event cells did not predict recognition memory success (Extended Data Fig. 11b, d; NB: *F* (1, 70) = 0.75, *p* = 0.39; SB: *F* (1, 70) = 1.1, *p* = 0.30; HB: *F* (1, 70) = 2.13, *p* = 0.15). Second, the strength of phase-locking of boundary cells neither predicted recognition memory success (Fig. 4b, d; NB: *F* (1, 82) = 1.16, *p* = 0.28; SB: *F* (1, 82) = 1.87, *p* = 0.18; HB: *F* (1, 82) = 0.45, *p* = 0.5) nor temporal order memory (Extended Data Fig. 10b, d; *F* (1, 82) = 1.33, *p* = 0.25; SB: *F* (1, 82) = 0.14, *p* = 0.71; HB: *F* (1, 82) = 1.98, *p* = 0.16).

In sum, boundary cells and event cells predict distinct aspects of memory formation: whereas the firing rate of boundary cells was predictive of later recognition memory performance, the phase-locking of event cells was predictive of temporal order memory performance.

### Neural state shifts across boundaries improve scene recognition but impair time discrimination

We next investigated the changes in the neural responses following boundaries at the population level of all n=580 MTL cells (pseudopopulation, see Methods). We examined the dynamics of population activity around the boundaries by evaluating the change of activity using principal component analysis. During NB video clips, the neural state exhibited only slow changes as a function of time (Fig. 5a, black dot marks the middle of the clip). In contrast, the neural state changed abruptly during SB and HB video clips following the boundaries (Fig. 5b, c; black dot marks boundary time). These abrupt ‘neural state shifts’ were consistent with the changes in firing rates we reported for boundary and event cells (but note here all recorded cells in the MTL were analyzed). To quantify the size of state shifts, we computed the multidimensional Euclidean distance MDD(*t*) in state space between a time *t* and the boundary (Fig. 5d-g; time of boundary is defined as *t* = 0 for convenience). The dimensionality of the state space we used was the number of PCs that together explained ≥ 99% of the variance. Plotting MDD as a function of time revealed an abrupt change within ~300 ms after the boundary for SB and HB video clips (Fig. 5d-g).

**Fig. 5:**
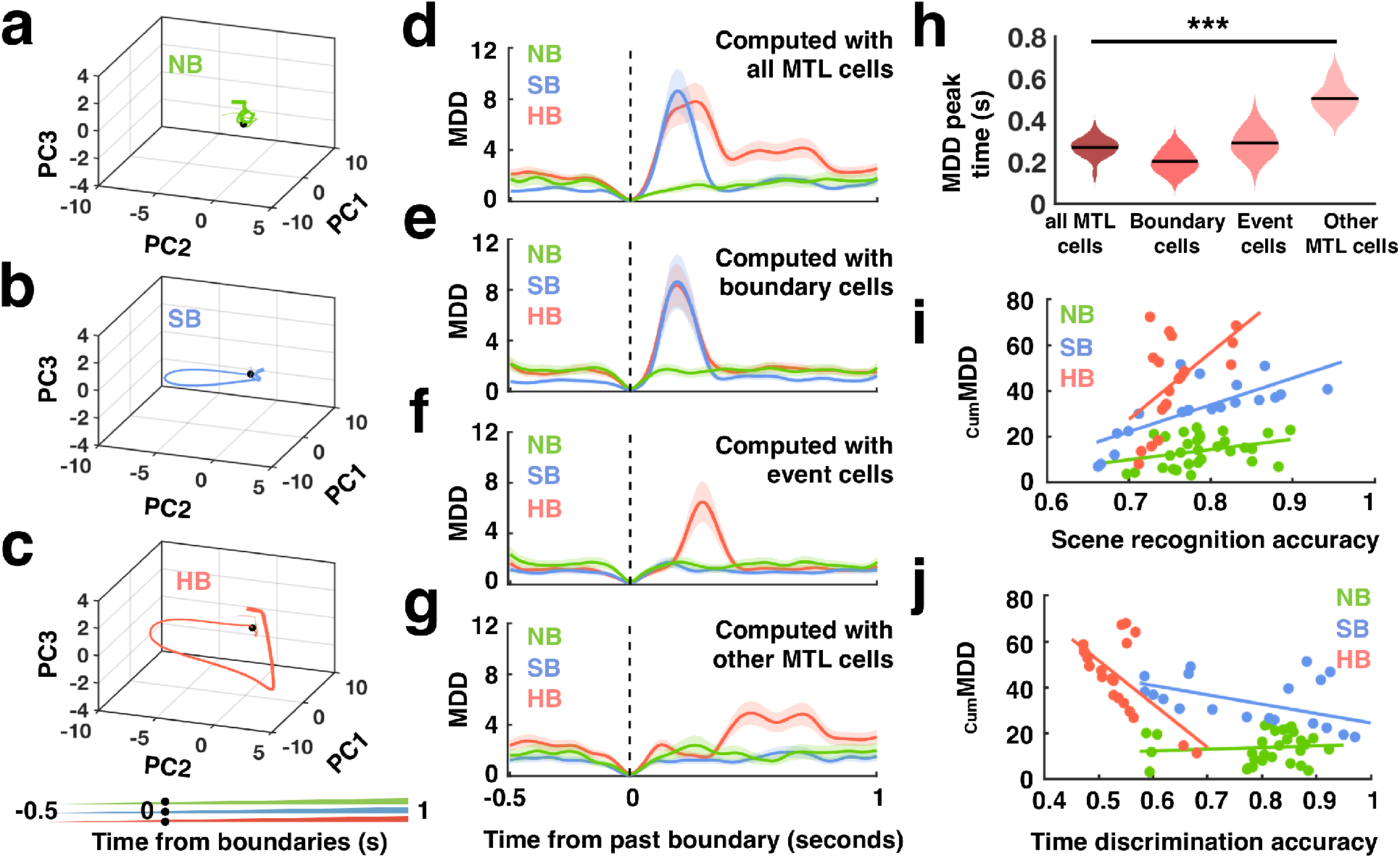
Population neural state shift magnitude following episodic transitions reflects subjects’ subsequent memory performance. **a-c.** Trajectories in neural state space formed by the top three principal components (PCs with most explained variance: PC1 = 26.05%, PC2 = 10.89%; PC3 = 6.69%) summarizing the activity of all MTL cells during the encoding stage for clips containing NB (**a**), SB (**b**) and HB (**c**). Each data point indicates the neural state at a specific time relative to boundary onset (line thickness indicates time; see scale on bottom). Black dots mark the time of the boundary (SB, HB) or the middle of the clip (NB). **d-g.** Multidimensional distance (MDD, i.e., Euclidean distance relative to boundaries in the PC space formed by all PCs that cover explained variance ≥ 99%, see **Extended Data Fig. 12**) as a function of time aligned to the middle of the clip (green: NB) and boundaries (blue: SB, red: HB). MDD is shown for all MTL cells (**d**, n = 528 in top 55 PCs space), all boundary cells (**e**, n = 42 in top 27 PCs space), all event cells (**f**, n = 36 in top 26 PCs space), and all other MTL cells (i.e., non-boundary/event cells in the MTL; **g**, n = 450 in top 58 PCs space). Shaded areas represent ± s.e.m. across trials. **h.** Latency analysis. Time when MDD shown in **d-g** reached peak value following HB (red lines) significantly differed when computed with different groups of cells. **i-j.** Correlation between distance in state space and behavior. **i.** Positive correlation between cumulative MDD (cumMDD: cumulative sum of Euclidean distances between boundary onset and the point of time at which the later target frame was shown in the PC space shown in **d**) and scene recognition accuracy. Dots mark the accuracy in the scene recognition task (x-axis) and the cumMDD during encoding (y-axis) of the target frames plotted separately for frames that follow NB (green: r = 0.33, p = 0.07), SB (blue: r = 0.75, p = 2×10^−4^) and HB (red: r = 0.54, p = 0.015). **j.** Negative Correlation between the cumMDD versus time discrimination accuracy plotted in the same format as **i** for NB (green: r = 0.097, p = 0.61), SB (blue: r = −0.49, p = 0.03) and HB (red: r = −0.64, p = 0.002). \*\*\**P* < 0.001, one-way ANOVA, degrees of freedom = (3, 76) in **h**.

We evaluated what types of cells contributed most to the neural state shift. First, neural state shifts following SBs were only visible when boundary cells were included (Fig. 5d, e). Second, early neural state shifts after HBs were only visible when event cells were included while later HB-related shifts remained present in the absence of either event cells (Fig. 5f) or both event and boundary cells (Fig. 5g). Third, the point of time at which MDD reached its maximal value varied systematically between groups of cells: the boundary responses carried by boundary cells appeared significantly earlier than those carried by boundary cells and non-boundary/event cells (Fig. 5h; *F* (3, 76) = 103.96, *p* = 8×10^−27^). Together, this shows that early population-level state shifts are principally due to the activity of boundary cells, whereas event cells and non-boundary/event cells in MTL contribute to slower-latency HB-related state shifts.

We next assessed whether the size of neural state shifts following boundaries during encoding were related to whether a stimulus was later remembered or not. We computed the extent and length of state changes in the population following a boundary by calculating the cumulative Euclidean distance traversed in state space in the period between the boundary and the tested frame (cumMDD; see Methods). We found that cumMDD was positively correlated with recognition accuracy for frames that followed both SB and HB, but not NB (Fig. 5i; Pearson correlation; SB: *r* = 0.75, *p* = 2×10^−4^; HB: *r* = 0.54, *p* = 0.015; NB: *r* = 0.33, *p* = 0.07). In contrast, cumMDD was negatively correlated with accuracy in the time discrimination task for both SB and HB, but not for NB (Fig. 5j, Pearson correlation, SB: *r* = −0.49, *p* = 0.03; HB: *r* = −0.64, *p* = 0.002; NB: *r* = 0.097, *p* = 0.61). Together, this result reveals a neural correlate of the tradeoff between these two types of memory, with large neural state shifts beneficial for recognition memory but detrimental for order memory. This observation is concordant with the behavioral results, where we found that HB-type boundaries enhanced recognition memory but disrupted order memory (Fig. 2).

### Reinstatement of neural context after boundaries facilitates recognition

It is thought that reinstatement of the neural context present at encoding enables mental time travel during memory retrieval^23,24^. However, it remains unknown what exactly is reinstated for continuous experience and how boundaries shape the retrieval process. To address this question, for each subject, we quantified the degree of reinstatement by computing the correlation between the vectors of spike counts of all recorded MTL neurons during recognition (1.5s fixed time window) and during encoding (1.5s sliding window, step size 0.1s; see Methods). Correct targets (i.e., frames from presented clips during encoding that were correctly remembered as “old”) were accompanied by a high correlation between neural activity during the scene recognition and the encoding period shortly after SB/HB transitions (Fig. 6a, e; p < 0.01, permutation test, see Methods). In contrast, we observed no significant correlation for forgotten targets (i.e., frames from presented clips that were incorrectly marked as “new”; Fig. 6b, f) or correctly identified foils (i.e., frames from unpresented clips correctly marked as “new”; Fig. 6c, g). Notably, correlations were strongest at points of time following the boundary that preceded the time at which the later tested boundary was shown (see Extended Data Fig. 13 for statistics). This observation indicates that what was reinstated was the neural state present following the boundary rather than the state present at the time the tested frame was shown.

**Fig. 6:**
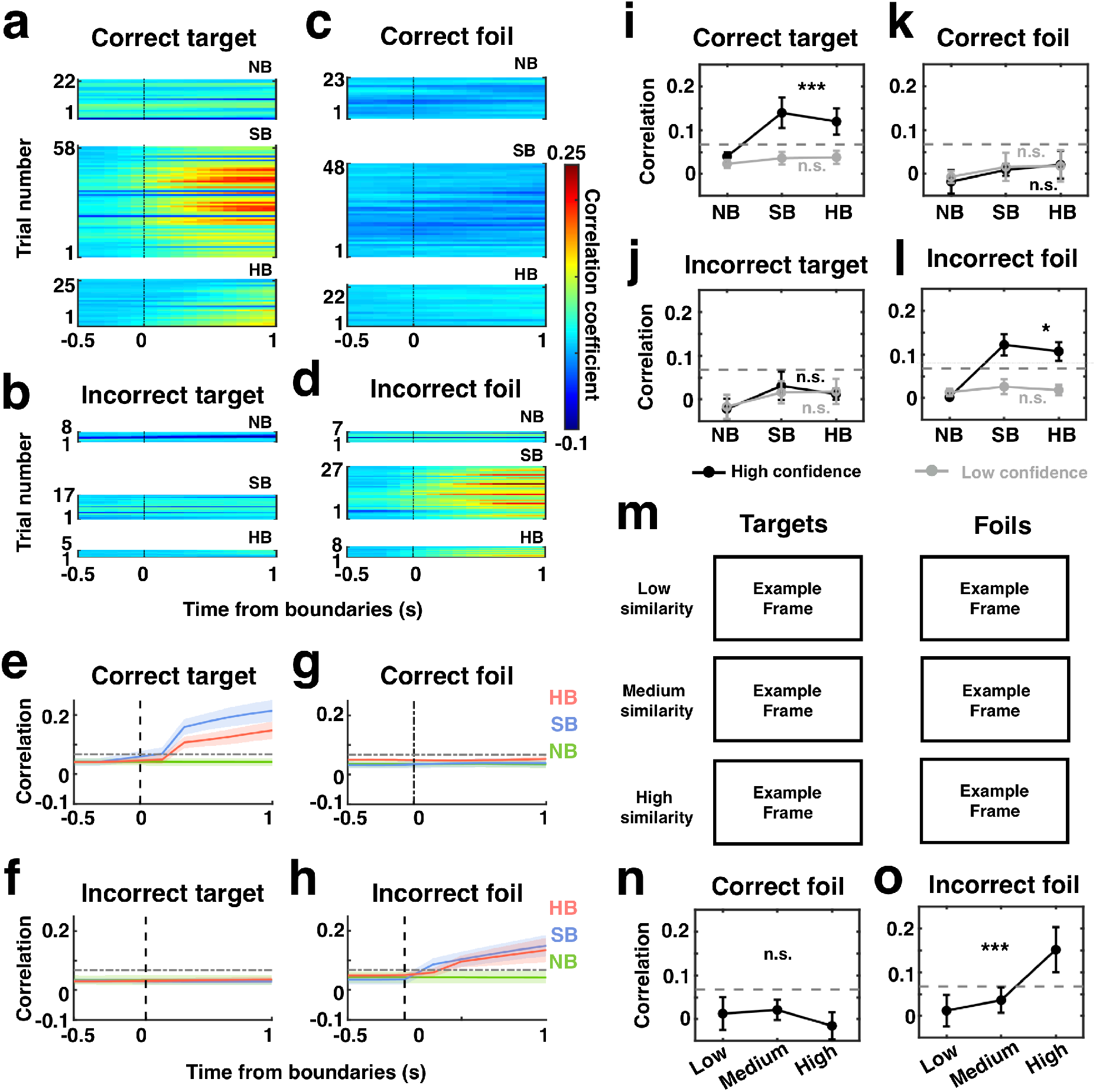
Reinstatement of neural context after boundaries during recognition. **a-d.** Single-subject example. Color code indicates correlation between the population response during scene recognition (0-1.5s relative to stimulus onset) and the encoding period (sliding window of 1.5s and 100ms step size). Correlations are aligned to the middle of the clip (NB) or boundaries (SB, HB) and are shown separately for correctly recognized familiar target (**a**), correctly recognized novel (not seen) foils (**c**), forgotten target (**b**) and incorrectly recognized foils (false positives, **d**) in the scene recognition task. **e-h.** Population summary. Correlation coefficient as shown in part **a-d**, averaged across all subjects for NB (green), SB (blue), and HB (red) trials. Shaded areas represented ± s.e.m. across subjects. The grey dashed horizontal lines denote the significant threshold (p < 0.01, permutation test, see Methods). **i-l.** Population summary (confidence). Reinstatement differed between frames remembered with high (black) and low (gray) confidence responses for “old” decisions (correct target and incorrect foil) but not ‘new” decisions (correct foil and incorrect target), regardless of whether they were correct or incorrect. Correlation coefficients as shown in part **e-h**, averaged over 0-1s after boundaries. **m.** Example target and foil frame pairs with low, medium and high similarity (rated by an independent control group, n = 30, see **Extended Data Fig. 4a)**. Note that owing to copyright issues, all original images have been removed but are available upon reasonable request. **n-o.** Population summary (target-foil similarity). Correlation coefficients versus similarity ratings between targets and foils, plotted for correct (**n**) and incorrect recognized foils (**o**). Error bars indicate standard deviation across n= 19 sessions. \**P* < 0.05, \*\*\**P* < 0.001, one-way ANOVA, degrees of freedom = (2, 54). Correlation metrices aligned to targets are shown in **Extended Data Fig.13**.

Reinstatement is thought to contribute primarily to recollection of details during retrieval^25^. Compatible with this view, we found that the correlations following boundary transitions were significantly stronger in high-compared to low-confidence trials during both correctly remembered as well as falsely recognized trials (Fig. 6i-l; High confidence, correct target: *F* (2, 54) = 8.61, *p* = 6×10^−4^; High confidence, incorrect foil: *F* (2, 54) = 5.76, *p* = 0.033).

Notably, strong correlations also occurred when a new image was incorrectly classified as seen before (Fig. 6d, h; incorrect foil; p < 0.01, permutation test, see Methods), thereby revealing a neural explanation for the false alarms. Were these false alarms, which were accompanied by neural reinstatement, caused by visual similarity between the targets and foils? To address this question, we assessed the similarity between each target and foil by acquiring similarity ratings from an independent control group of subjects (n = 30). Similarity values were balanced across NB, SB and HB (As shown in Extended Data Fig. 4a). We split foils into low (top 66.67% - 100%), medium (top 33.33% - 66.67%) and high (top 1% – 33.33%) similarity groups (see examples in Fig. 6m). Correlations between encoding and scene recognition were significantly stronger for highly similar foils falsely recognized as old compared to low and medium similarity foils (Fig. 6o; incorrect foil: *F* (2, 54) = 10.67, *p* = 1×10^−4^). In contrast, the extent of correlation for correctly rejected foils (true negatives) did not vary as a function of similarity (Fig. 6n, correct foil: *F* (2, 54) = 2.182, *p* = 0.144). This result indicates that the reason for false alarms was that the wrong context was reinstated due to the high similarity of the foil with a target. Together, these results support the notion that the neural state present at encoding following the boundary was reinstated during memory retrieval.

## Discussion

Memories are often conceptualized as discrete events on a narrative timeline^5^. However, the very definition of an event is only beginning to be understood. Specifically, where do events start and end, and how are multiple signals bound together over time to form a singular event? Here, we tested the hypothesis that boundary detection is a mechanism that segments continuous experience into discrete events by causing “jumps” in the neural context. Behaviorally, consistent with previous findings^26^, we found that boundaries enhanced recognition memory while disrupting temporal order memory. Neuronally, we observed single neurons in the medial temporal lobe (MTL) that signaled the presence of boundaries with increased firing rates. These cells triggered a neural state shift at the population level. Aspects of the boundary-related responses predict specific kinds of later improved (recognition) or impaired (order memory) memory performance, showing the behavioral relevance of this neural mechanism for shaping our memories. Lastly, successful recognition relied on the reinstatement of the neural state present shortly after a detected boundary during retrieval.

Both soft-and hard boundaries are accompanied by salient visual changes, whereas the ‘no boundary’ control condition had no such changes (Fig. 1b). However, the observed responses to boundaries of boundary cells and event cells (Fig. 3) cannot be explained by these sharp visual input changes: First, the cells did not respond to the equally strong visual changes at stimulus onsets and offsets (Extended Data Fig. 7–8). Second, cells differentiated between soft-and hard boundaries despite these types of boundaries not being distinguishable at the pixel-and higher-level visual features (we also confirmed this finding psychophysically with additional human control subjects, Extended Data Table 1).

Boundary cells respond to both soft-and hard boundaries, whereas event cells respond only to hard boundaries (Fig. 3). The distinct responses of these two kinds of cells might reflect the hierarchical structure of episodic memory, with event cells representing episodic transitions between completely distinct events while boundary cells represent more frequent but smaller episodic transitions within the same overall narrative. Our findings provide empirical evidence for the theory^8^ that event segmentation is a hierarchical dynamic procedure, with fine to coarse segmentations associated with different kinds of cognitive boundaries. The anatomical location and response latency of our cells was also compatible with this proposal: boundary cells (which responded first) were most common in the parahippocampal gyrus, whereas event cells (which responded later) were most common in the hippocampus (Extended Data Table 4). Thus, the faster more frequently occurring boundary-related responses were encoded in parahippocampal gyrus, with more abstract and rarer boundary-related responses encoded later in the hippocampus. This distinction was also visible at the population level, with neural state shifts for SBs mainly driven by boundary cells, whereas HB-related state shifts occurred later and were driven by a broader group of cells (Fig. 5d-h). We hypothesize that the early responses of boundary cells (mostly in the parahippocampal gyrus) reflect contextual changes detected in the higher-level visual areas^27,28^, while event cells (mostly in the hippocampus) are the result of a late output signal from a comparator operation^29,30^ (between predicted and received signal).

The responses of boundary or event cells in our study bring to mind border and place cells in the rodent hippocampus^14,31^, which respond to physical boundaries or locations in the environment, respectively. As rodents move between compartments, place cells cluster at boundaries (e.g., doorways)^12^, crossing of which is followed by remapping^16,17^ or reinstatement^13,18^ of a different set of hippocampal place fields. In our study, boundary cells and event cells responded to transitions (boundaries) between different episodic contexts in the video clips with an increase in firing rates. Also, similar to place field remapping, neural state in the medial temporal lobe changed abruptly following crossing of a boundary. Notably, this occurred despite the new context entered being novel (a situation in which place fields would take multiple exposures to develop). When subjects were re-exposed to familiar target frames during the later recognition test, reinstatement of neural state occurred (similar to remapping) if the item was successfully recognized. Similar to place field reinstatement, the reinstated neural state was most similar to that which occurred at the point of time shortly following the boundary that preceded the tested memory item rather than the neural state that was present when the tested frame was first shown. This finding provides insight into the important question of what neural context seen at what point of time in the past is reinstated during mental time travel and memory search^32–36^. This finding also indicates that abrupt changes in neural context are important to demarcate periods of time that can be reinstated later from those that cannot. We note several key differences between the boundary cells and event cells we reported here, and border cells reported in rodents. Border cells respond to physical boundaries and rely on specific tasks in specific physical environments in which rodents are extensively trained. In contrast, boundary or event cells in humans responded flexibly to abstract cognitive boundaries for a large variety of different contexts (videos), none of which subjects have seen before and in each of which the boundary is marked in a different way in a different narrative. This property is an essential requirement for a process to divide experience into episodes to shape episodic memory, which by definition each occur only once in novel environments.

What roles do the boundary response play in episodic memory formation and retrieval? At the single-cell level, responses of boundary cells and event cells during encoding were predictive of different aspects of subjects’ subsequent memory performance. Whereas the firing rate of boundary cells predict recognition memory strength, the phase-locking of event cells to theta oscillations predicted temporal order memory accuracy. This indicates that the two kinds of cells played distinct roles, with each strengthening only one kind of memory using a different plasticity mechanism (firing rate and phase-locking, respectively). At the population level, the strength and duration of the neural state shift that followed a boundary was positively correlated with the strength of recognition memory and negatively correlated with the strength of temporal order memory. This opposing effect of neural state shifts provides a neural explanation for a fundamental trade-off imposed by segmenting memory into discreet episodes: such an approach to organize memory weakens temporal order memory but enhances memory for items seen shortly after initiating a new episode^1,6^. Our subjects also exhibited this tradeoff behaviorally. Together, our results provide direct neural evidence for representations of cognitive boundary detections and the role of these signals in initiating the formation of and structuring neural representations of episodic memory.

## Methods

### Task

The task consisted of three parts: encoding, scene recognition, and time discrimination (Fig. 1a).

#### Encoding

subjects watched a series of 90 unique clips with no sound and were instructed to memorize as much of the clips as possible. Each trial started with a baseline period (i.e., a fixation cross at the center of a blank screen; the duration of the baseline period ranged from 0.9s to 1.1s across all the trials). The fixation period was followed by the presentation of a video clip that contained either no boundaries (NB, continuous movie shots), soft boundaries (SB, cuts to a new scene within the same movie, 1 to 3 SB per clip, randomly distributed in the clips), or a hard boundary (HB, cuts to a new scene from a different movie, 1 HB per clip at 4 seconds of the clip) Examples of SB and HB are shown in Fig. 1b and Extended Data Fig. 1. A yes/no question related to the content of the clip (e.g., Is anyone in the clip wearing glasses?) appeared randomly after every 4-8 clips.

#### Scene recognition

After watching all 90 clips, subjects’ memory for the content of the videos was evaluated in a scene recognition test. During scene recognition, frames extracted from encoded clips (target frames), and frames from new, never shown, clips (foil frames) were presented to the subjects. Subjects were instructed to identify whether these frames were “old” or “new” (i.e., whether they had seen the frame during the encoding session). Two target frames (in total n = 180) were extracted from each clip, one randomly pulled out from the first half of the clip and the other one from the second half. Then 50% of these extracted target frames from both first/second half of the clip (n = 90) were selected as templates to search for foil frames from a different movie played by different actors/actresses (n = 30), a different movie played by the same actors/actresses (n = 30), or the unpresented portion from the same movie played by same actors/actress (n = 30) to introduce different levels of similarity between the target frames and foil frames. The total number of target/foil frames (n = 30 targets and 30 foils for each boundary type) and the average similarity level of foil frames were counterbalanced across different boundary types (F (1, 88) = 2.62, p = 0.11; rated by Amazon Mechanical Turk workers, see Methods, *Similarity ratings*).

#### Time discrimination

After the scene recognition test, we evaluated subjects’ memory about the temporal structure of the clip with a time discrimination test. In each trial, two frames (half of them picked at 1s and 7s, and the other half picked at 3s and 5s of the clip) separated by different kinds of boundaries (NB, SB, or HB) were extracted from the encoded clips and were presented side by side. Subjects were instructed to indicate which of the two frames (i.e., “left” or “right”) appeared first (earlier in time) in the videos they watched during encoding.

#### Confidence measurement

All binary choices were made together with a subjective confidence judgment (i.e., sure, less sure, very unsure). Thus, there were always 6 possible responses for each question.

### Similarity ratings

#### Visual properties of SB and HB

Both SB and HB transitions were accompanied by transient visual changes (cuts to a scene), whereas there were no such visual changes for the control NB condition. We quantified the visual changes of each boundary type by computing metrics that relate to pixel level differences: Luminance; Contrast; Complexity; Entropy; Color distribution) between pre- and post-boundary frames. In addition, to quantify visual differences not directly captured at the pixel level, we used pre- and post-boundary frames as inputs for AlexNet network (pretrained on ImageNet dataset)^37^, extracted the activation matrices from the layer ‘fc7’ for both images and computed the Euclidean distance between their activation matrices. Moreover, we collected perceptual ratings (i.e., similarity ratings between pre- and post-boundary frames) from Amazon Mechanical Turk (MTurk) workers. During similarity ratings, pre- and post-boundary frames were presented side by side and MTurk workers were instructed to rate the similarity between them by clicking on a scaling bar ([0 1]; 0 = different, 1 = identical). See results in Extended Data Table 1.

#### Similarity ratings between target and foil frames

When selecting foil frames, we used target frames as templates to search for foil frames with different similarity levels (see Methods, *Task*). We presented the target frame with its corresponding foil frame side by side and instructed MTurk workers to rate the similarity between them (see results in Extended Data Fig. 4a).

#### Time discrimination without encoding

To ensure the time discrimination task could not be solved by pure reasoning, we recruited MTurk workers to perform the time discrimination test without watching clips (see results in Extended Data Fig. 4b).

### Subjects

#### Patients

Twenty patients (see patients’ demographics in Extended Data Table 3) with drug-resistant epilepsy volunteered for this study and gave their informed consents. The institutional review boards of Toronto Western Hospital and Cedars-Sinai Medical Center approved all protocols. The task was conducted while the patients stayed in the hospital after implantation of depth electrodes for monitoring seizures. The location of the implanted electrodes was solely determined by clinical needs. The behavioral analyses included results from all 20 subjects and the neural results were analyzed across 19 subjects (Subject #20 had no usable recordings, see patient information in Extended Data Table 3).

#### Amazon Mechanical Turk Workers (MTurk workers)

MTurk workers were recruited for similarity ratings (see Methods, *Similarity ratings*), including 30 subjects (age 23.25 ± 3.42, 9 female) for rating the visual properties of different boundaries (Extended Data Table. 1), 30 subjects (age 22.79 ± 5.73, 11 female) for rating the similarity between target and foil frames (Extended Data Fig. 4a), and 30 subjects (age 25.06 ± 6.11, 7 female) for performing the time discrimination task without encoding session (Extended Data Fig. 4b). All control tasks conducted on Amazon Mechanical Turk were under the approval of the institutional review board of Boston Children’s Hospital and informed consents were obtained with digital signatures for each subject.

### Electrophysiology

We recorded bilaterally from the amygdala, hippocampus, and parahippocampal gyrus, as well as other regions outside the medial temporal lobe using hybrid depth electrodes (Ad-Tech company, Oak Creek, Wisconsin, USA), which contained eight 40-μm diameter microwires at the tip of each electrode shank. For each microwire, broadband signals (0.1-9000 Hz filtered) were recorded at 32 kHz using the ATLAS system (Neuralynx Inc., Bozeman, Montana, USA).

### Spike sorting and quality metrics of single units

The recorded signals were filtered offline in the 300 to 3000 Hz band, with a zero-phase lag filter. Spikes were detected and sorted using the semiautomated template-matching algorithm Osort^38,39^. We computed spike sorting quality metrics for all putative single units (see Extended Data Fig. 6) to quantify our recording and sorting quality^40–42^.

### Electrode localization

Electrode localization was based on postoperative MRI/CT scans. We co-registered postoperative and preoperative MRIs using Freesurfer’s mri_robust_register^43^. To summarize electrode positions and to provide across-study comparability, we aligned each subject’s preoperative scan to the CIT168 template brain in MNI152 coordinates^44^ using a concatenation of an affine transformation followed by a symmetric image normalization (SyN) diffeomorphic transform^45^. The MNI coordinates of the 8 microwires from the same electrode shank were marked as one location. MNI coordinates of microwires with putative neurons detected across all the subjects were plotted on a template brain for illustration (Fig. 1e).

### Data analyses

#### Boundary cell

For each recorded neuron, we counted spikes within the [0 1] seconds (post boundary) and [−1 0] seconds time interval (baseline) relative to boundaries during encoding. A cell was considered a boundary cell if it met the following criteria: 1) its spike counts within post boundary time windows were significantly different from its spike count within baseline time windows for SB and HB but not for NB (p < 0.05, permutation t-test); 2) its spike counts within post boundary time windows were significantly greater in SB and HB than NB (p < 0.05, permutation t-test).

#### Event cell

A cell was considered as an event cell if it met the following criteria: 1) its spike counts within post boundary time windows were significantly different from its spike count within baseline time windows for HB but not for NB and SB (p < 0.05, permutation t-test); 2) its spike counts within post boundary time windows were significantly greater in HB than NB and SB (p < 0.05, permutation t-test).

#### Boundary and event cells’ responses to stimulus onsets and offsets

For each selected boundary cell and event cells, we counted spikes within the [0 1]s (post) and [−1 0]s (pre) time interval relative to stimulus onsets/offsets during encoding. The boundary cell or event cell was considered as not responding to stimulus onsets/offsets if their spike counts did not differ between post and pre window in all three boundary conditions (p > 0.05, permutation t-test).

### Chance level for cell response analyses

To estimate the number of neurons that would be considered boundary cells or event cells by chance, we repeated the same procedures for boundary cell and event cell analyses after randomly shuffling the trial labels (NB, SB, HB) 1000 times. For each iteration, we obtained the proportion of selected boundary cells and event cells relative to the total number of neurons within each region. These 1000 values formed the empirically estimated null distribution for the proportion of boundary cells and event cells expected by chance. A region was considered to have a significant amount of boundary cells or event cells if its actual fraction of significant cells exceeded 95% of the null distribution (Extended Data Table 4; p < 0.05).

### Association between spiking activity during encoding and later memory retrieval accuracy

#### Firing rate modulation

For each boundary cell and event cell, we grouped its spike activity within [0 1] seconds after boundaries during encoding based on subjects’ subsequent memory performance either in the scene recognition task (correct versus incorrect recognition) or the time discrimination task (correct versus incorrect discrimination). We then computed the firing rate as a function of time (bin size = 200ms, step size = 2ms) for each trial, which was further z-score normalized using the mean and standard deviation of the firing rate across the whole trial. For each boundary cell and event cell, we then computed the mean z-scored firing rate within [0 1] seconds time interval relative to boundaries for each trial and averaged this value across trials within each boundary type. The resulting values across all boundary cells and event cells were used for comparisons across NB, SB and HB trials (Figure. 4c, g and Extended Data Fig. 10c and 11c).

#### Phase modulation

For each spike of each boundary cell and event cell, we computed the phase of the spike relative to the theta-frequency band filtered local field potential signals (LFP) recorded from the same microwire. The original LFP signals were first band-passed between 1-300Hz and downsampled from 32Khz to 500Hz. We performed automatic artifact rejections^46^ accompanied with manual visual inspections (ft_databrowser.m from Fieldtrip toolbox^47^) to remove large transient signal changes (e.g., spike-artifacts and interictal discharges) from the downsampled LFPs. Next, we extracted theta-band oscillatory activity (4-8Hz) from the artifact-rejected signals using the BOSC method^48^, which detects periods of transient oscillatory activity (disregarding periods of time with non-sinusoidal signals). For each detected bout of theta, we then applied a Hilbert transform to obtain theta phase as a function of time and extracted the phase for each spike fired by boundary and event cells. The phase locking strength of each boundary or event cell was quantified as the mean resultant length (MRL) of all spike phases of all spikes fired within [0 1] seconds after boundaries (0 = no phase locking; 1 = strongest phase locking) with theta bouts detected. The resulting MRL values were then compared across NB, SB and HB trials between conditions (Figure. 4d, h and Extended Data Fig. 10d and 11d). The computation of MRL is sensitive to sample number. Therefore, the comparison of MRL between correct and incorrect trials was conducted with balanced spike counts.

### State-space analyses

#### Neural state trajectories

For each trial, we binned each neuron’s spike counts during encoding into non-overlapping 10-ms wide bins, followed by smoothing with a 200ms standard deviation Gaussian kernel and z-score normalization (mean and standard deviation were calculated across the entire trial). We used these z-scored smoothed spike density estimates from all recorded MTL cells across all subjects to form a pseudocopulation. We applied principal component analysis (PCA) to reduce the dimensionality of the pseudopopulation (MATLAB function svd.m). We then rank-ordered the resulting principal components (PCs) by their explained variance (function dpca_explainedVariance.m from dpca toolbox^49^) and plotted the average neural state trajectories for each boundary type in a three-dimensional space constructed by the top three PC components (Fig. 5a-c).

#### Multidimensional distance (MDD)

MDD was defined as the Euclidean distance between two points in the PC space (with PCs accounted for > 99% explained variance; see Extended Data Fig. 12).

#### Cumulative Multidimensional distance (cumMDD)

cumMDD was defined as the cumulative sum of all Euclidean distance values between two points of time (in the PC space).

### Reinstatement of neural context

This analysis was done separately for each session of simultaneously recorded neurons and did not rely on the pseudopopulations defined in the previous section.

#### Correlation between encoding and retrieval

Neural activity was quantified for each neuron in bins of 1.5s width and a step size of 100ms. We computed the Pearson correlation coefficients (corrcoef.m from MATLAB) between the neural population activity during scene recognition (1 time bin x number of neurons) and encoding (80 time bins x number of neurons) at each time step.

#### Significant correlation threshold

We computed the same correlation values after randomly shuffling the trial labels (i.e., after disrupting the correspondence between encoding and scene recognition trials) to obtain the average correlation strength across trials and neurons expected by chance. This procedure was repeated 1000 times to form a null distribution, in which the 2.5^th^ and 97.5^th^ percentile values were used as the threshold to determine significance of the actual correlation values (dashed horizontal lines in Fig. 6).

#### Comparison between boundary-aligned and target/foil aligned correlation

We computed the average correlation coefficients within [−1 0] and [0 1] seconds relative to boundary or relative to the time when target frames present in the original clip. We then defined boundary-aligned/target aligned correlations by subtracting the average correlation coefficients in [−1 0] from [0 1] seconds relative to boundary/target, respectively. Notably, for foil frames, we used the time when their corresponding target frames (see Methods, *Task*) appeared in the original clip for alignment.

### Statistics

For comparisons between two groups, we used the permuted t-test statistic, and for comparisons between more than two groups, we used parametric F-statistics. For statistical thresholding, permutation tests were conducted to generate a null distribution estimated from 1000 runs on data with scrambled labels, which avoids the assumption of normality when evaluating significance.

## Data and Code availability

Data and analytic code that support the findings of this study will be deposited at Open Science Framework upon acceptance.

## Acknowledgements

We thank Mengmi Zhang, Jan Kaminski and other members from Rutishauser and Kreiman lab for discussion; Nand Chandravadia and Victoria Barkely for data transferring and organization; Chrystal Reed, Jeffrey Chung, and the clinical teams at both Cedars-Sinai Medical Center and Toronto Western Hospital. We especially indebted to the patient volunteers who participated in this study. This work was supported by the Brain Canada program (to A.GPS.) and NIH U01NS103792 (to U.R.).

## Author contributions

J.Z. conceived the project. J.Z., G.K. and U.R. contributed ideas for experiments and analysis. V.T. and A.M. managed patients and surgeries; J.Z., A.GPS., M. Y. and C. M. collected data; J.Z. performed the analyses; J.Z., G.K. and U.R. wrote the manuscript with input from all authors.

---

**Extended Data Table. 1:** Comparison of visual attributes among different types of boundaries. The visual difference between pre boundary frames and post boundary frames (i.e., B_post_ - B_pre_) was quantified for each property: 1) Luminance: average pixel value of the grayscale image; 2) Contrast: standard deviation across all pixels of the grayscale image^50^;3) Complexity: JPEG size of an image with a compression quality setting of 80 (on a scale from 1 to 1000)^51^. Simple images are highly compressible, resulting in smaller file size; 4) Entropy: as an additional index of image complexity. It is computed from the histogram distribution of the 8-bit gray-level intensity values. Entropy varies with the “randomness” of the image, with low entropy associated with less complex images; 5) Color distribution: each picture was converted to the CIE L*A*B color space, which approximates characteristics of the human visual system^52^. L dimension corresponds to luminance, A and B dimension corresponds to two chromatic channels ranging from red to green, and from blue to yellow, respectively. 6) Higher-level features: the higher-level features of images were quantified as the activations from the layer ‘fc7’ from an AlexNet network^37^ trained on the ImageNet data set. The difference of higher-level features was computed as the Euclidean distance between the activation matrices obtained from the post boundary frame and the pre boundary frame. 7) Similarity: the similarity between the post boundary frame and the pre boundary frame was rated by an independent group of Amazon Mechanical Turk workers (n = 30). The rating scale was 0 to 1 (0 = totally different, 1 = identical). The statistical significance was evaluated using one-way ANOVA (Analysis of Variance) test with post-hoc Tukey HSD (Honestly Significant Difference) test.

| Visual attributes | $B_{\text{post}} - B_{\text{pre}}$ | | | Boundary Type main effect | Post-hoc test | | |
| --- | --- | --- | --- | --- | --- | --- | --- |
|  | NB | SB | HB |  | NB vs SB | NB vs HB | SB vs HB |
| Luminance difference | $4.47 \pm 0.23$ | $4.95 \pm 0.53$ | $5.18 \pm 0.78$ | $P < 0.001$ | $P < 0.01$ | $P < 0.001$ | $P = 0.27$ |
| Contrast difference | $1.25 \pm 0.19$ | $1.93 \pm 0.50$ | $1.77 \pm 0.27$ | $P < 0.001$ | $P < 0.001$ | $P < 0.001$ | $P = 0.18$ |
| Complexity difference | $498.38 \pm 1.04$ | $1971.38 \pm 130.03$ | $2031.09 \pm 236.23$ | $P < 0.001$ | $P < 0.001$ | $P < 0.001$ | $P = 0.30$ |
| Entropy difference | $0.19 \pm 0.01$ | $0.43 \pm 0.06$ | $0.46 \pm 0.08$ | $P < 0.001$ | $P < 0.01$ | $P < 0.001$ | $P = 0.12$ |
| LABL difference | $0.33 \pm 0.03$ | $0.69 \pm 0.05$ | $0.68 \pm 0.16$ | $P < 0.001$ | $P < 0.001$ | $P < 0.001$ | $P = 0.42$ |
| LABA difference | $0.08 \pm 0.01$ | $0.14 \pm 0.08$ | $0.14 \pm 0.06$ | $P < 0.001$ | $P < 0.001$ | $P < 0.001$ | $P = 0.53$ |
| LABB difference | $0.16 \pm 0.08$ | $0.34 \pm 0.22$ | $0.36 \pm 0.24$ | $P < 0.001$ | $P < 0.01$ | $P < 0.001$ | $P = 0.19$ |
| Alexnet fc7 Euclidean distance | $22.18 \pm 1.98$ | $217.24 \pm 27.80$ | $199.77 \pm 53.85$ | $P < 0.001$ | $P < 0.001$ | $P < 0.001$ | $P = 0.14$ |
| Similarity | $1.00 \pm 0.00$ | $0.67 \pm 0.13$ | $0.63 \pm 0.15$ | $P < 0.001$ | $P < 0.001$ | $P < 0.001$ | $P = 0.37$ |

**Extended Data Table. 2:** MNI coordinates for electrodes plotted in Figure 1e.

| Electrode ID | PatientID | Location | Side | MNI coordinates |  |  |
| --- | --- | --- | --- | --- | --- | --- |
|  |  |  |  | X | Y | Z |
| 1 | TWH101 | Hippocampus | Left | -23.09 | -14.45 | -18.87 |
| 2 | P64CS | Hippocampus | Left | -22.56 | -5.84 | -16.74 |
| 3 | P60CS | Hippocampus | Left | -30.57 | -16.02 | -14.95 |
| 4 | TWH101 | Hippocampus | Left | -28.98 | -18.92 | -14.12 |
| 5 | TWH103 | Hippocampus | Left | -23.01 | -17.72 | -13.23 |
| 6 | TWH109 | Hippocampus | Left | -21.22 | -18.65 | -11.97 |
| 7 | TWH111 | Hippocampus | Left | -26.04 | -20.12 | -15.33 |
| 8 | TWH116 | Hippocampus | Left | -27.21 | -17.88 | -14.73 |
| 9 | TWH117 | Hippocampus | Left | -30.84 | -20.08 | -11.07 |
| 10 | TWH125 | Hippocampus | Left | -20.18 | -18.98 | -19.96 |
| 11 | TWH126 | Hippocampus | Right | 25.59 | -3.43 | -27.55 |
| 12 | TWH127 | Hippocampus | Right | 24.94 | -13.49 | -23.44 |
| 13 | P61CS | Hippocampus | Right | 30.18 | -10.78 | -18.39 |
| 14 | P62CS | Hippocampus | Right | 34.7 | -26.87 | -6.36 |
| 15 | TWH110 | Hippocampus | Right | 24.74 | -17.67 | -13.86 |
| 16 | TWH111 | Hippocampus | Right | 26.44 | -17.83 | -14.01 |
| 17 | TWH113 | Hippocampus | Right | 27.77 | -24.86 | -14.92 |
| 18 | TWH138 | Hippocampus | Right | 27.01 | -26.45 | -11.18 |
| 19 | TWH125 | Amygdala | Left | -20.49 | -3.57 | -16.21 |
| 20 | P64CS | Amygdala | Left | -21.67 | -5.32 | -16.45 |
| 21 | P65CS | Amygdala | Left | -19.19 | -2.97 | -19.11 |
| 22 | TWH103 | Amygdala | Left | -16.59 | -4.88 | -20.67 |
| 23 | TWH127 | Amygdala | Left | -20.01 | -1.72 | -21.18 |
| 24 | TWH129 | Amygdala | Left | -17.96 | -5.21 | -18.36 |
| 25 | P63CS | Amygdala | Right | 20.57 | -5.94 | -24.84 |
| 26 | P60CS | Amygdala | Right | 24.75 | -1.87 | -20.98 |
| 27 | TWH113 | Amygdala | Right | 16.82 | -3.86 | -22.04 |
| 28 | TWH117 | Amygdala | Right | 17.43 | -5.71 | -17.96 |
| 29 | TWH129 | Amygdala | Right | 20.21 | -4.88 | -11.76 |
| 30 | TWH120 | Amygdala | Right | 19.01 | -1.82 | -16.13 |
| 31 | P61CS | Parahippocampus | Left | -25.81 | -22.86 | -8.64 |
| 32 | TWH109 | Parahippocampus | Left | -31.01 | -27.01 | -14.12 |
| 33 | TWH113 | Parahippocampus | Left | -35.77 | -20.97 | -9.12 |
| 34 | TWH116 | Parahippocampus | Left | -30.72 | -22.83 | -9.25 |
| 35 | TWH120 | Parahippocampus | Left | -35.79 | -20.99 | -9.11 |
| 36 | TWH125 | Parahippocampus | Left | -30.72 | -22.83 | -9.12 |
| 37 | TWH127 | Parahippocampus | Right | 26.22 | -34.57 | -9.98 |
| 38 | TWH129 | Parahippocampus | Right | 25.88 | -35.62 | -10.03 |
| 39 | TWH138 | Parahippocampus | Right | 27.11 | -36.63 | -10.57 |

**Extended Data Table 3.** Subject information.

| Patient ID | Subject ID | Age | Gender | Number of detected cells |  |  |  |  |
| --- | --- | --- | --- | --- | --- | --- | --- | --- |
|  |  |  |  | All cells (985) | Boundary cell in MTL (42) | Boundary cell outside MTL (8) | Event cell in MTL (36) | Event cell outside MTL (9) |
| P60CS | 1 | 67 | Male | 70 | 1 | 1 | 2 | 0 |
| P61CS | 2 | 52 | Female | 74 | 2 | 0 | 3 | 0 |
| TWH101 | 3 | 25 | Female | 50 | 1 | 0 | 0 | 1 |
| THW103 | 4 | 49 | Male | 57 | 4 | 1 | 0 | 1 |
| TWH106 | 5 | 26 | Male | 42 | 0 | 0 | 0 | 0 |
| TWH109 | 6 | 28 | Male | 49 | 3 | 1 | 2 | 1 |
| TWH110 | 7 | 38 | Male | 31 | 1 | 0 | 1 | 0 |
| TWH111 | 8 | 63 | Female | 55 | 3 | 1 | 1 | 2 |
| TWH113 | 9 | 36 | Male | 54 | 3 | 1 | 3 | 1 |
| P62CS | 10 | 25 | Female | 51 | 3 | 0 | 2 | 1 |
| TWH116 | 11 | 25 | Female | 44 | 2 | 0 | 1 | 0 |
| TWH117 | 12 | 68 | Female | 21 | 2 | 0 | 0 | 0 |
| P64CS | 13 | 63 | Female | 70 | 3 | 1 | 3 | 0 |
| P65CS | 19 | 55 | Female | 78 | 4 | 0 | 5 | 0 |
| TWH120 | 15 | 25 | Female | 49 | 2 | 0 | 2 | 1 |
| TWH125 | 15 | 24 | Male | 51 | 3 | 1 | 2 | 1 |
| TWH126 | 16 | 25 | Male | 22 | 0 | 0 | 1 | 0 |
| TWH127 | 17 | 28 | Male | 44 | 2 | 1 | 2 | 0 |
| TWH129 | 18 | 25 | Female | 47 | 3 | 0 | 4 | 0 |
| TWH138 | 19 | 46 | Female | 26 | 0 | 0 | 2 | 0 |
| TWH100 | 20 | 20 | Female | Psychophysics only, No single neuron recording |  |  |  |  |

**Extended Data Table. 4:** Distribution of boundary and event cells across brain areas. Shown is the proportion of all recorded cells that qualified as boundary cells and event cells (AMY: amygdala; HPC: hippocampus; PHG: parahippocampal gyrus; OFC: orbitofrontal; ACC: anterior cingulate cortex; MCC: middle cingulate cortex; SMA: supplementary motor area; INS: insula) and the significance of this proportion against the null distribution (see Methods). Significant entries are marked in gray.

| Cell Type | AMY | HPC | PHG | OFC | ACC | MCC | SMA | INS |
| --- | --- | --- | --- | --- | --- | --- | --- | --- |
| Boundary cell | 7.10%<br>(12/169)<br>p = 0.009 | 3.50%<br>(12/343)<br>p = 0.036 | 26.47%<br>(18/68)<br>p = 0.001 | 1.59%<br>(3/126)<br>p = 0.158 | 2.13%<br>(2/94)<br>p = 0.103 | 0.00%<br>(0/32)<br>p = 0.638 | 1.45%<br>(1/69)<br>p = 0.232 | 2.38%<br>(2/84)<br>p = 0.089 |
| Event cell | 4.27%<br>(7/169)<br>p = 0.011 | 7.87%<br>(27/343)<br>p = 0.004 | 2.94%<br>(2/68)<br>p = 0.041 | 6.35%<br>(6/126)<br>p = 0.021 | 2.13%<br>(2/94)<br>p = 0.103 | 0.00%<br>(0/32)<br>p = 0.638 | 0.00%<br>(0/69)<br>p = 0.638 | 1.19%<br>(1/84)<br>p = 0.277 |

**Extended Data Table. 5:** ANOVA test with the reinstatement of neural context at boundaries and targets, Related to Figure 6. Statistical numbers (i.e., F-numbers and p-values) for correlation values (as shown in Fig. 6e-h, averaged within [0,1] seconds relative to boundaries or when target presented in the clip) covaried with independent variables (Confidence: high, medium, low; BoundaryType: NB, SB, HB; Time distance: relative distances between targets and boundaries in [0,1], [1,2], [2,3], [3,4] seconds; Similarity: similarity ratings between foils and their corresponding targets) and their interaction terms. Significant main effects or interactions are marked in gray. \**P* < 0.05, \*\**P* < 0.01, \*\*\**P* < 0.001.

| Corect Target vs Encoding<br>Correlation Coefficient | Boundary aligned |  | Target aligned |  |
| --- | --- | --- | --- | --- |
|  | F-number | P-value | F-number | P-value |
| Confidence | 2.56 | > 0.05 | 1.01 | > 0.05 |
| BoundaryType | **5.77** | **0.007\*\*** | 0.82 | > 0.05 |
| Time distance | 2.04 | > 0.05 | 1.33 | > 0.05 |
| Confidence * BoundaryType | **9.77** | **0.0009\*\*\*** | 0.88 | > 0.05 |
| Confidence * Time distance | 3.11 | > 0.05 | 1.11 | > 0.05 |
| BoundaryType * Time distance | 2.88 | > 0.05 | 0.56 | > 0.05 |
| Confidence * Distance * BoundaryType | **6.43** | **0.013\*** | 1.24 | > 0.05 |

| Incorrect Target vs Encoding<br>Correlation Coefficient | Boundary aligned |  | Target aligned |  |
| --- | --- | --- | --- | --- |
|  | F-number | P-value | F-number | P-value |
| Confidence | 1.69 | > 0.05 | 0.66 | > 0.05 |
| BoundaryType | 2.78 | > 0.05 | 0.42 | > 0.05 |
| Time distance | 1.84 | > 0.05 | 1.34 | > 0.05 |
| Confidence * BoundaryType | 2.21 | > 0.05 | 0.77 | > 0.05 |
| Confidence * Time distance | 1.19 | > 0.05 | 1.08 | > 0.05 |
| BoundaryType * Time distance | 2.03 | > 0.05 | 0.59 | > 0.05 |
| Confidence * Distance * BoundaryType | 1.55 | > 0.05 | 1.53 | > 0.05 |

| Correct Foil vs Encoding<br>Correlation Coefficient | Boundary aligned |  | Target aligned |  |
| --- | --- | --- | --- | --- |
|  | F-number | P-value | F-number | P-value |
| Confidence | 2.26 | > 0.05 | 0.92 | > 0.05 |
| Boundary Type | 2.79 | > 0.05 | 0.77 | > 0.05 |
| Similarity | 2.48 | > 0.05 | 1.69 | > 0.05 |
| Confidence * BoundaryType | 1.12 | > 0.05 | 0.64 | > 0.05 |
| Confidence * Similarity | 1.78 | > 0.05 | 0.69 | > 0.05 |
| BoundaryType * Similarity | 1.43 | > 0.05 | 0.64 | > 0.05 |
| Confidence * Similarity * BoundaryType | 2.09 | > 0.05 | 1.44 | > 0.05 |

| Incorrect Foil vs Encoding<br>Correlation Coefficient | Boundary aligned |  | Target aligned |  |
| --- | --- | --- | --- | --- |
|  | F-number | P-value | F-number | P-value |
| Confidence | 2.47 | > 0.05 | 0.89 | > 0.05 |
| Boundary Type | **5.12** | **0.011\*** | 0.69 | > 0.05 |
| Similarity | **6.69** | **0.004\*\*** | 1.97 | > 0.05 |
| Confidence * BoundaryType | **7.12** | **0.003\*\*** | 0.72 | > 0.05 |
| Confidence * Similarity | 2.11 | > 0.05 | 0.65 | > 0.05 |
| BoundaryType * Similarity | 1.33 | > 0.05 | 0.74 | > 0.05 |
| Confidence * Similarity * BoundaryType | **9.19** | **0.0021\*\*** | 1.37 | > 0.05 |

**Extended Data Fig. 1:**
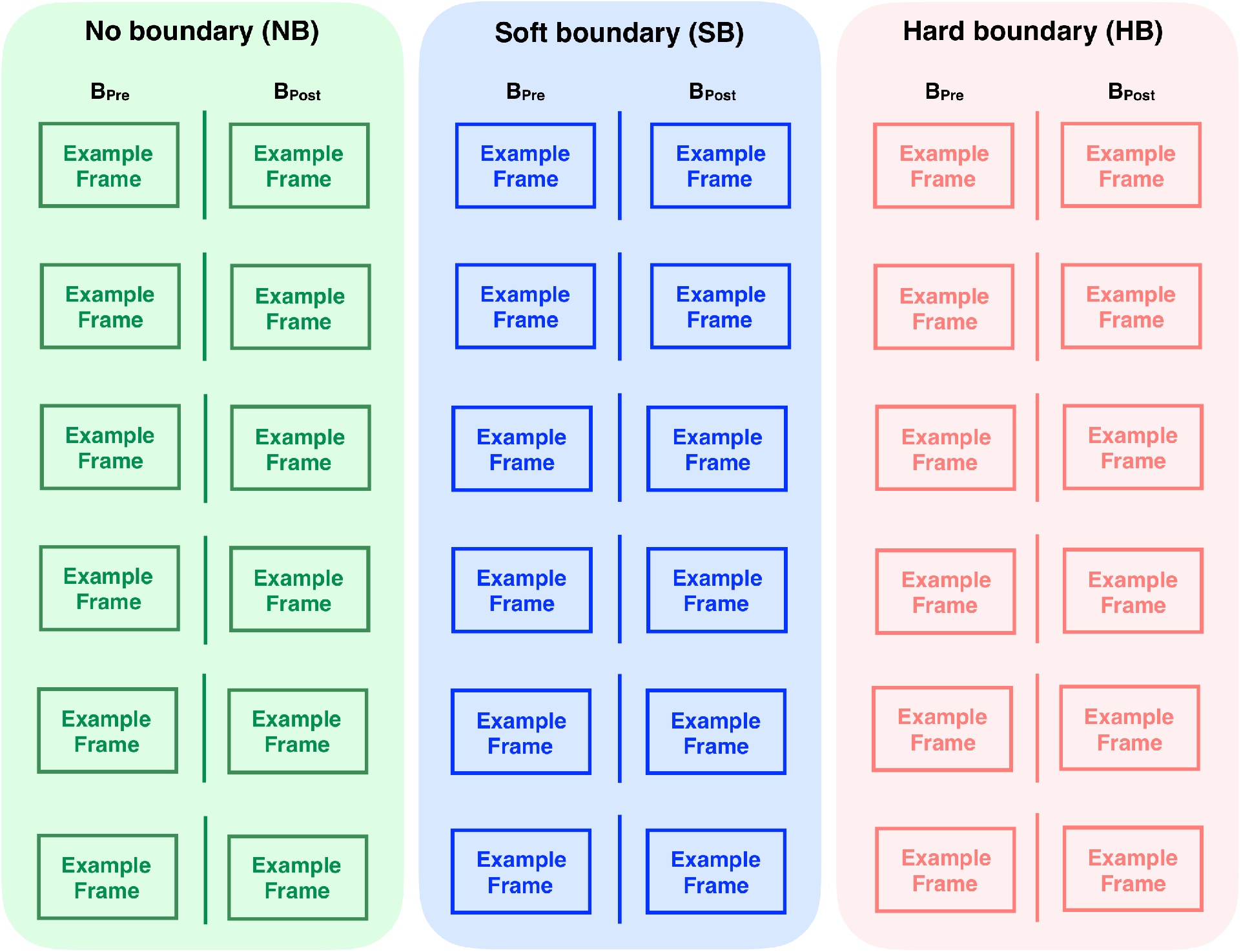
Examples of the three different types of boundaries: Six examples showing the frame before (pre) and after (post) the middle of the clip (NB, green) or the frame before and after soft boundaries (SB, blue, cuts between different shots of the same movie), or hard boundaries (HB, red, cut between shots from different movies). Note that owing to copyright issues, all original images have been removed but available upon reasonable request.

**Extended Data Fig. 2:**
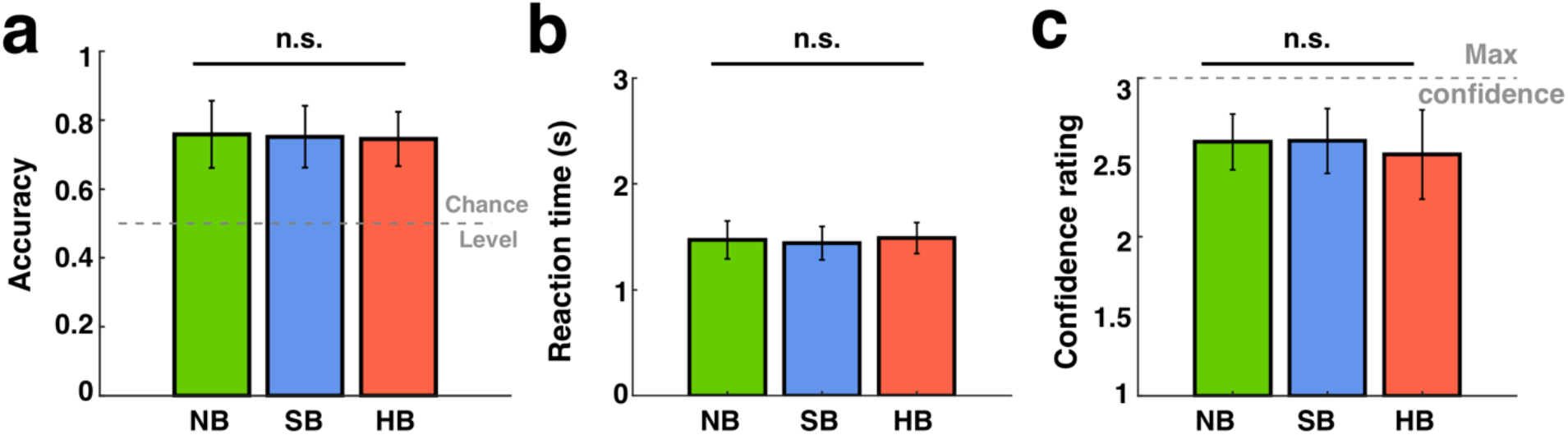
Subjects’ performance in the scene recognition task did not differ significantly across different boundary types. **(a-c)** Behavior quantified by accuracy (**a**), response time (**b**), and confidence level (**c**) across all trials. Results are shown for boundary type NB (green), SB (blue), and HB (red) during the scene recognition task. The horizontal dashed lines in (**a)** show chance levels (0.5) and in (**c)** show the maximum possible confidence value (3=high confidence). Error bars indicate standard deviation across n = 20 sessions. One-way ANOVA between NB/SB/HB, degrees of freedom = (2, 57).

**Extended Data Fig. 3:**
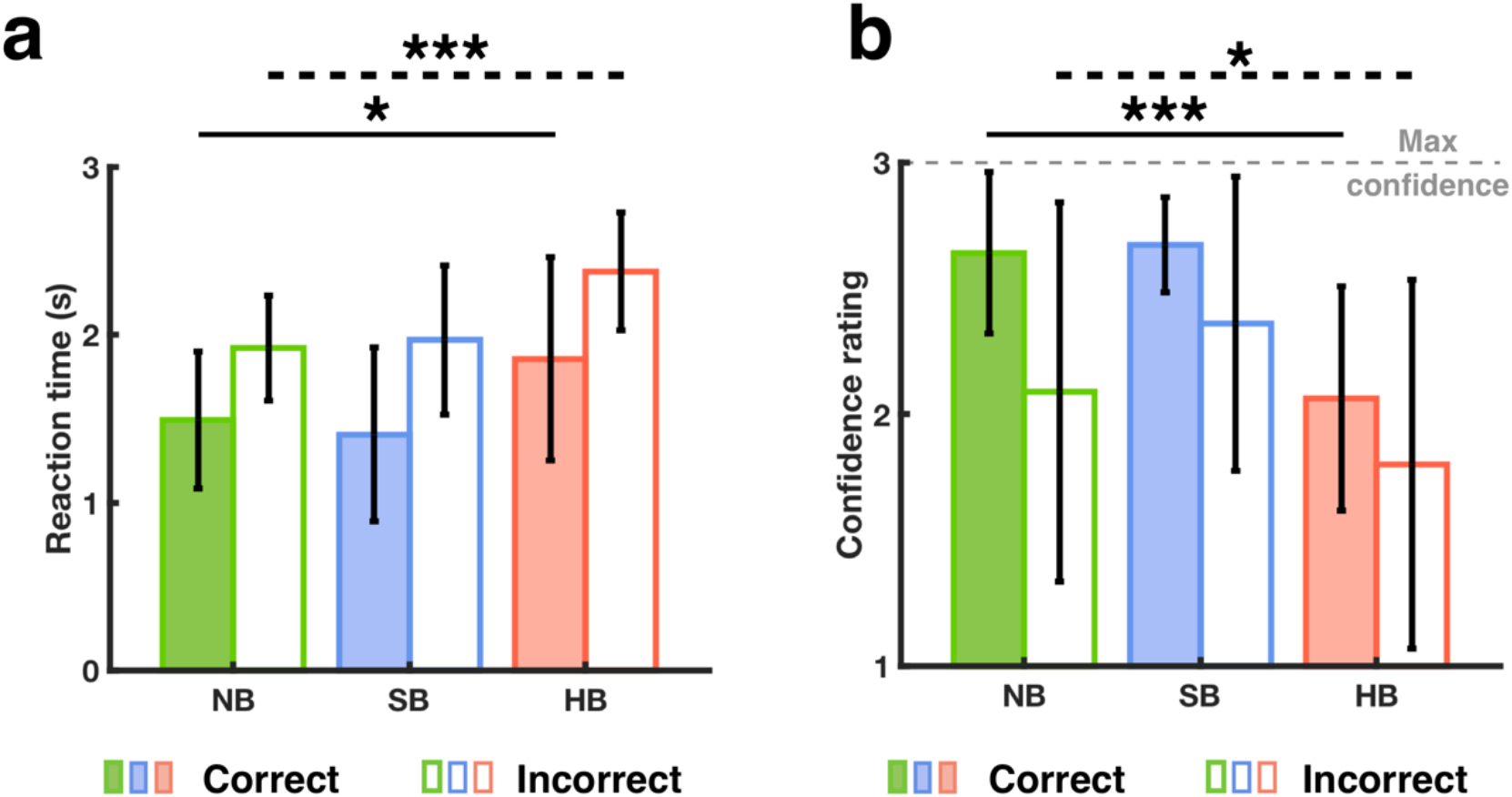
Longer reaction time and lower confidence level for HB compared to SB and NB, regardless of whether the clips’ temporal orders were remembered or forgotten. **(a-b)** Behavior quantified by response time (**a**) and confidence level (**b**) during the time discrimination task for clips whose temporal order were remembered (color filled) vs. forgotten (empty). Results are shown for boundary type NB (green), SB (blue), and HB (red). The horizontal dashed line in (**b)** show the maximum possible confidence value (3 = high confidence). For both correct (color filled) and incorrect trials (empty), subjects showed longer reaction times (**a**; Correct trials: HB = 1.86 ± 0.61 seconds, NB = 1.49 ± 0.41 seconds, SB = 1.40 ± 0.52 seconds, *F* (2, 57) = 4.29, *p* = 0.02; Incorrect trials: 2.38 ± 0.35 seconds, NB = 1.92 ± 0.31 seconds, SB = 1.96 ± 0.44 seconds, *F* (2, 57) = 9.06, *p* = 3.84×10^−4^) and lower confidence ratings (**b**; Correct trials: HB = 2.06 ± 0.45, NB = 2.64 ± 0.32, SB = 2.67 ± 0.19, *F* (2, 57) = 21.02, *p* = 1.45×10^−7^; Incorrect trials: HB= 1.80 ± 0.73, NB = 2.09 ± 0.75, SB = 2.35 ± 0.58, *F* (2, 57) = 3.23, *p* = 0.04) when discriminating between two frames earlier separated by a HB compared to SB and NB. Error bars indicate standard deviation across n = 20 subjects. \**P* < 0.05, \*\*\**P* < 0.001, one-way ANOVA between NB/SB/HB, degrees of freedom = (2, 57).

**Extended Data Fig. 4:**
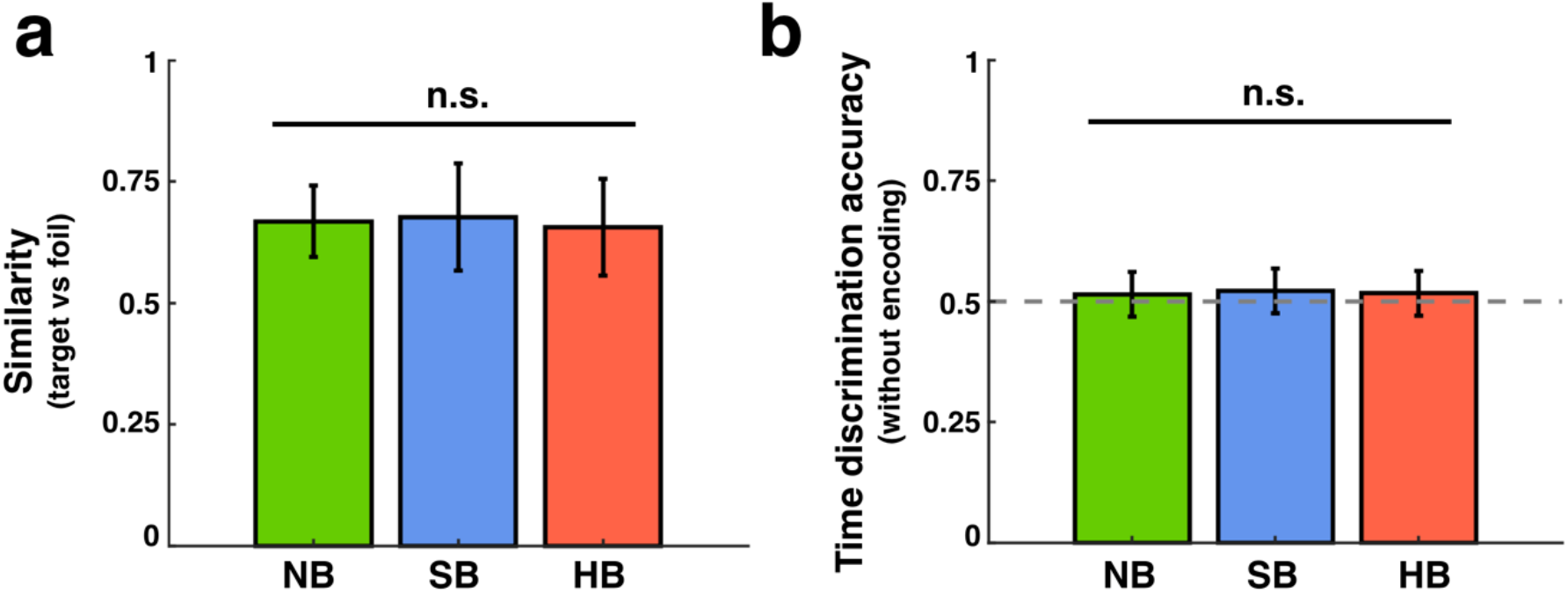
Boundary effect on time discrimination was not driven by the difficulty of scene recognition and pure reasoning. **a**, Similarity between corresponding target and foil frames used in the scene recognition task, ranging from 0 (totally different) to 1 (identical) as rated by an independent group of Amazon Mechanical Turk workers (n = 30). **b.** Time discrimination accuracy by an independent group of Amazon Mechanical Turk workers (n = 30) who did not watch the clips. Error bars indicate standard deviation across n=30 subjects. One-way ANOVA, degrees of freedom = (2, 87).

**Extended Data Fig. 5:**
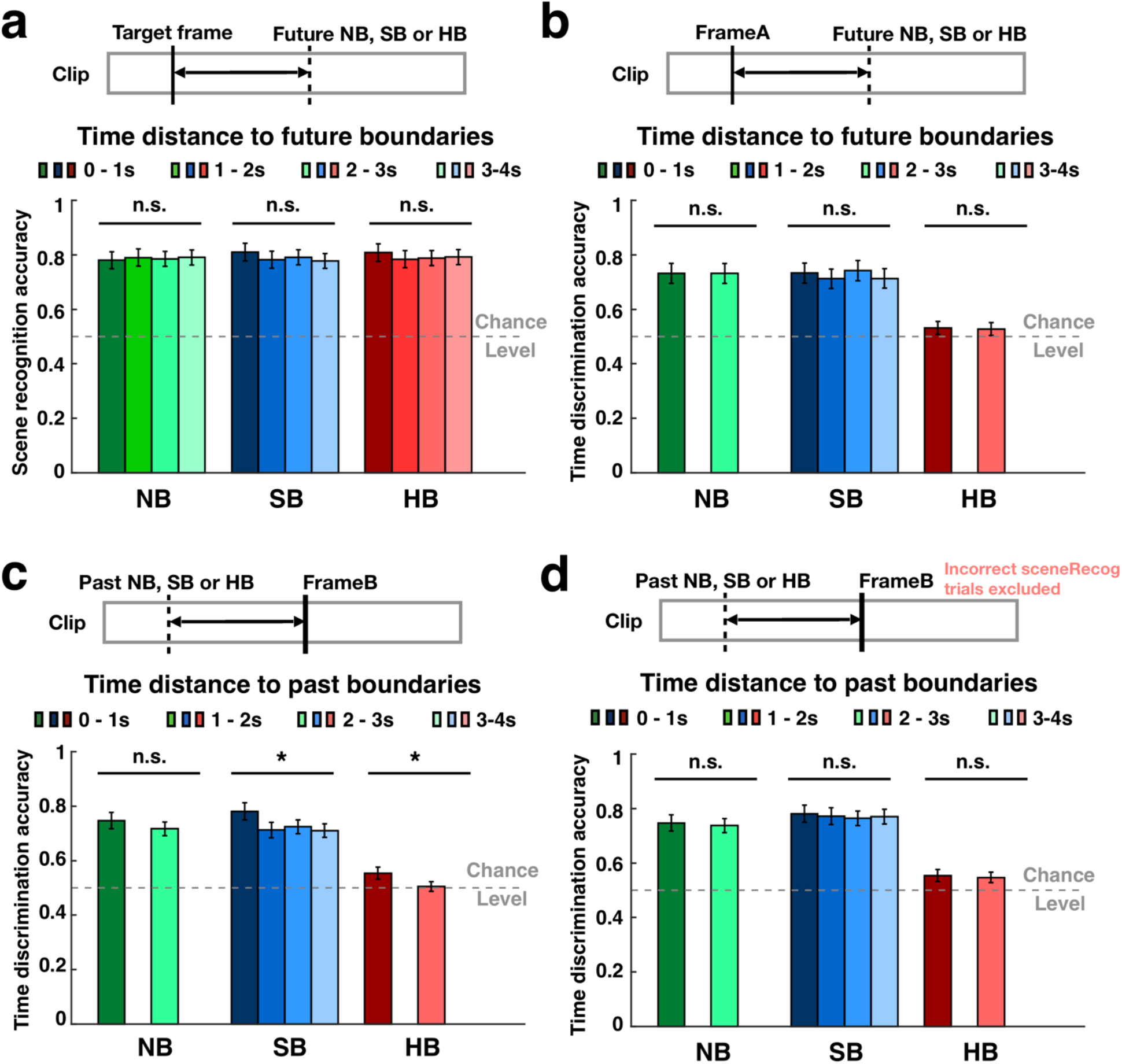
Scene recognition and time discrimination accuracy were not modulated by the distance between target and future boundaries: **a.** Expanding on the results in **Fig. 2d**, scene recognition accuracy for target frames grouped by the time elapsed between the target and the *future* boundary (note that **Fig. 2d** showed time elapsed from the *past* boundary). **b.** time discrimination accuracy grouped by the time elapsed between the target frame A and the *future* boundary. **c.** time discrimination accuracy grouped by the time elapsed between the target frame B and the *past* boundary. **d.** time discrimination accuracy grouped by the time elapsed between the target frame A and the *past* boundary *but excluding clips with incorrect scene recognition*. In **b-d**, there were no tested frames within [1,2] or [3,4] seconds from the NB and HB clips. Error bars indicate standard deviation across n = 20 subjects. \**P* < 0.05, one-way ANOVA, degrees of freedom = (3, 76) in **a**; degree of freedom = (1, 38) in **b-d** for NB and HB and degrees of freedom = (3, 76) for SB.

**Extended Data Fig. 6:**
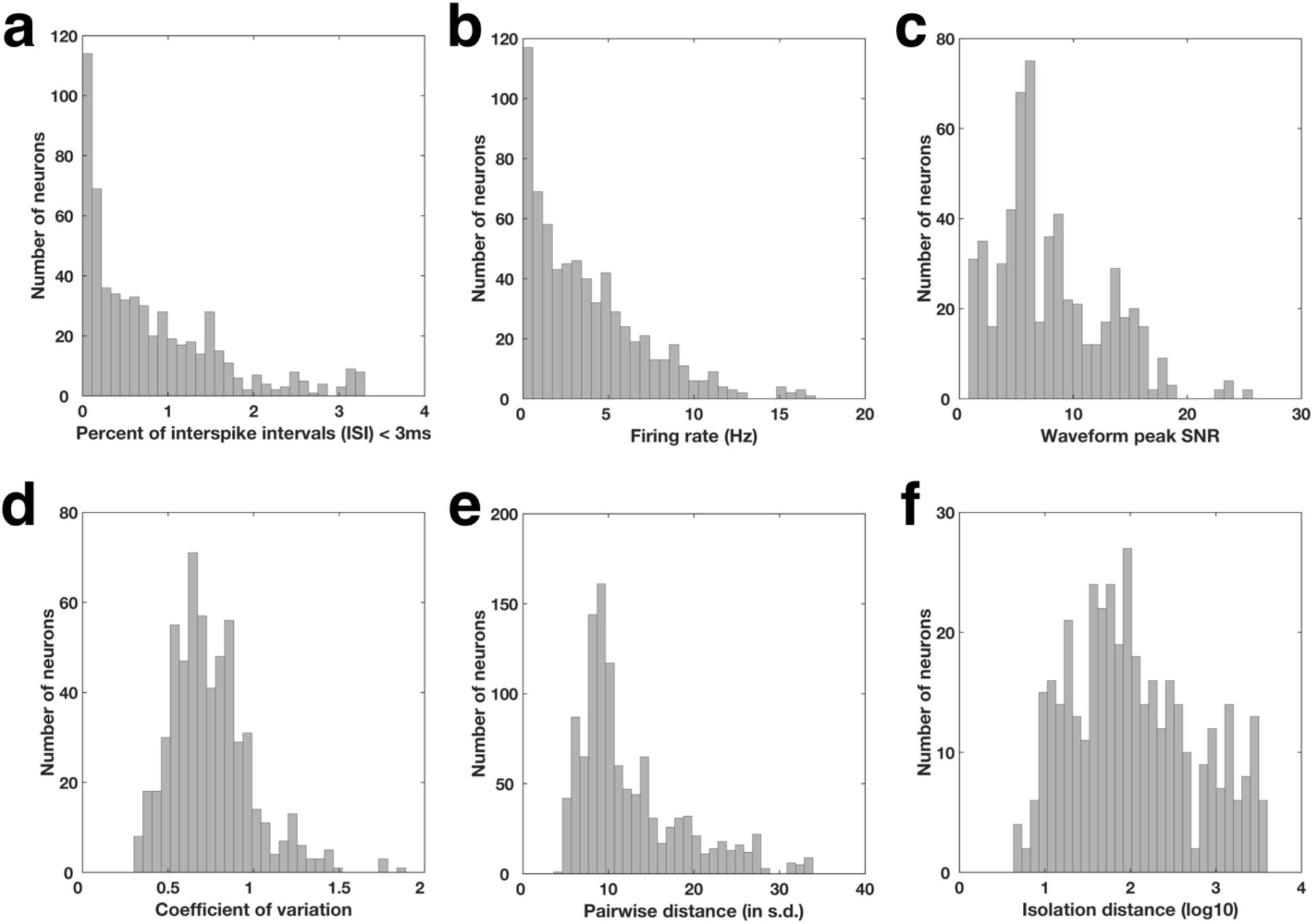
Spike sorting quality metrics for all identified putative single cells: **a**, Histogram of proportion of inter-spike intervals (ISI) that were shorter than 3ms (0.80% ± 0.80%, mean ± s.d.). **b**, Histogram of average firing rate within the entire recording session for all identified putative single cells (3.74 ± 3.34 Hz, mean ± s.d.). **c**, Histogram of waveform peak signal-to-noise ratio (SNR), which is the ratio between the peak amplitude of the mean waveform and the s.d. of the noise of each identified putative single cell (8.00 ± 4,73, mean ± s.d.). **d**, Histogram of coefficient-of-variation (CV2) in the ISI for each identified putative single cell (0.74 ± 0.24, mean ± s.d.). **e**, Histogram of the pairwise isolation distance between putative single cells identified from the same wire (projection test; 12.44 ± 6.18 s.d. of the signal,). **f**, Histogram of isolation distance across all identified putative single cells that was calculated in a ten-dimensional feature space of the energy normalized waveforms^42^. These quality metrics are comparable to previous published works in the field^40,41,53^.

**Extended Data Fig. 7:**
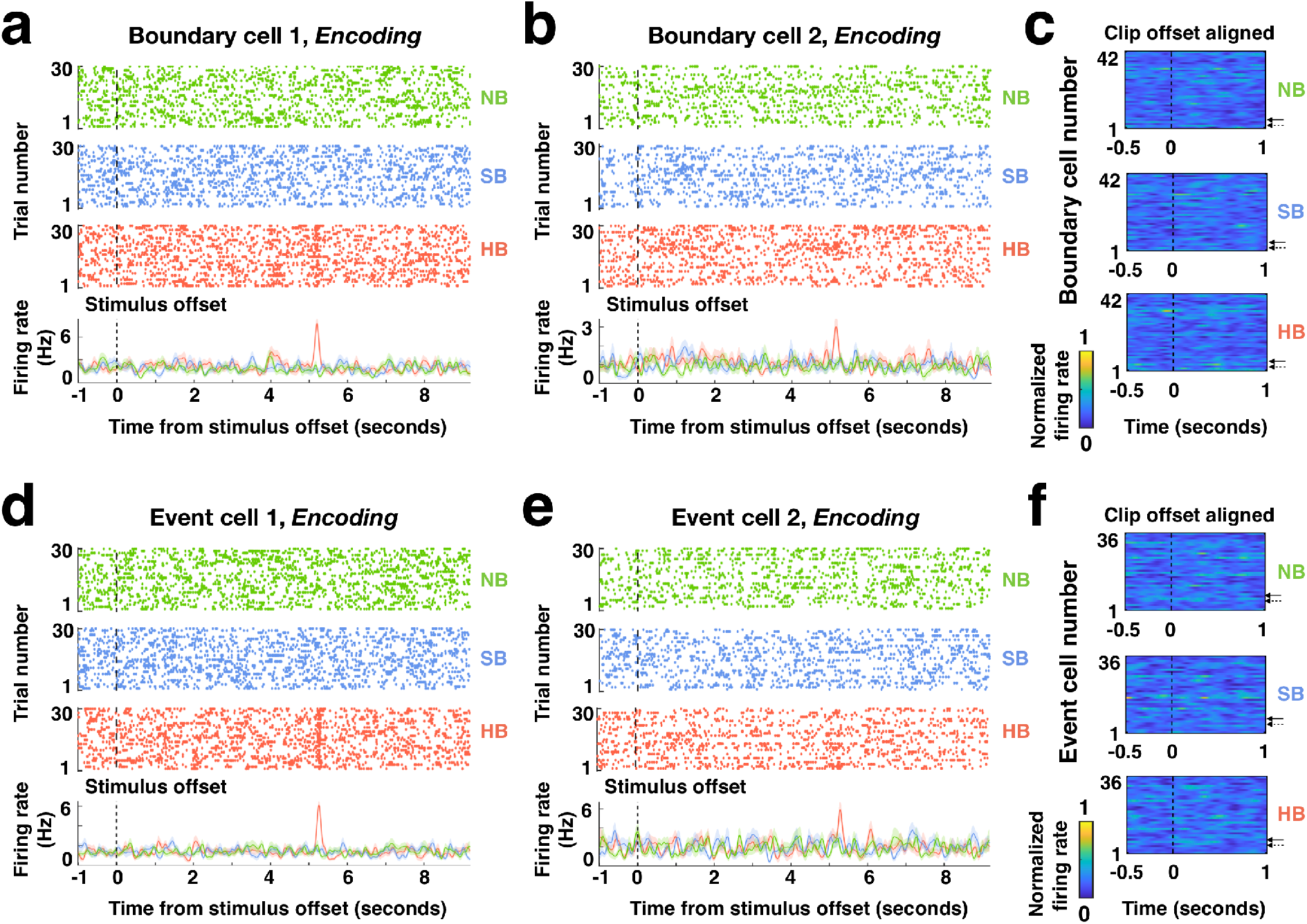
Boundary cells and event cells do not respond to clip offsets: **a-b**, Responses during the encoding stage from the same example boundary cells shown in **Fig.3a-b** aligned to the clip offsets. **c**, Firing rates of all 42 boundary cells (solid and dashed arrows denote the examples in **a** and **b**, respectively) during the encoding stage aligned to the clip offsets, averaged over trials within each boundary type and normalized to each neuron’s maximum firing rate throughout the entire task (see color scale on bottom). **d-e**, Responses during the encoding stage from the same example boundary cells shown in **Fig.3e-f** aligned to the clip offsets. Same format as **a-b**. **f**, Firing rates of all 36 event cells (solid and dashed arrows denote the examples in **d** and **e**, respectively) during the encoding stage, using the same format as **c**. For **a, b, d, e**, Top: raster plot color coded for different boundary types (green: NB; blue: SB; red: HB). Bottom: Post-stimulus time histogram (bin size = 200ms, step size = 2ms, shaded areas represented ± s.e.m. across trials). Black dashed lines indicate clip offsets.

**Extended Data Fig. 8:**
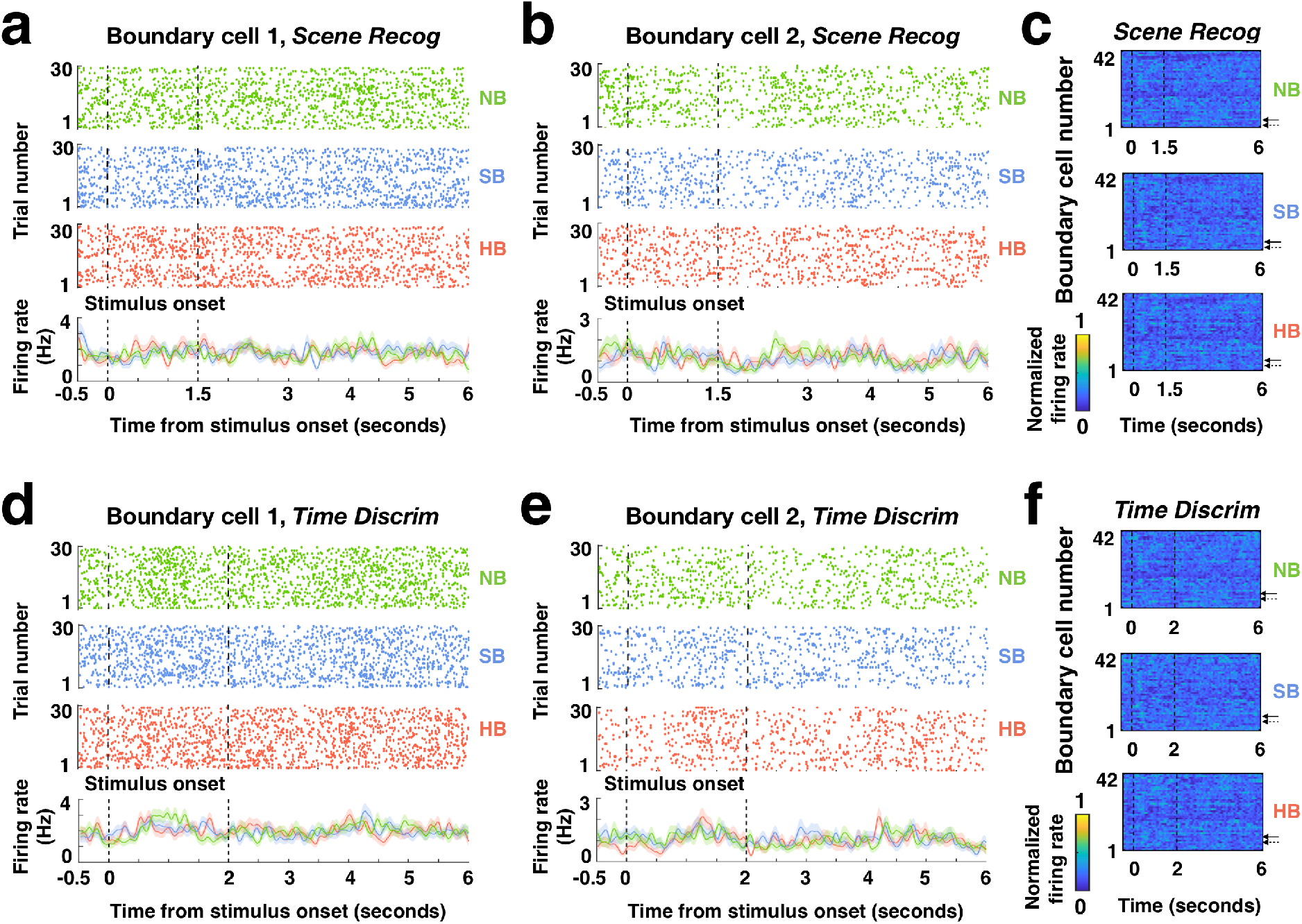
Responses of boundary cells during scene recognition and time discrimination: **a-b**, Responses during the scene recognition stage from the same example boundary cells shown in **Fig.3a-b** aligned to the stimulus onsets. **c**, Firing rates of all 42 boundary cells (solid and dashed arrows denote the examples in **a** and **b**, respectively) during the scene recognition aligned to the stimulus onsets, averaged over trials within each boundary type and normalized to each neuron’s maximum firing rate throughout the entire task (see color scale on bottom). **d-e**, Responses during the time discrimination stage from the same example boundary cells shown in **Fig.3e-f** aligned to the stimulus onsets. Same format as **a-b**. **f**, Firing rates of all 42 boundary cells (solid and dashed arrows denote the examples in **d** and **e**, respectively) during the time discrimination stage, using the same format as **c**. For **a, b, d, e**, Top: raster plot color coded for different boundary types (green: NB; blue: SB; red: HB). Bottom: Post-stimulus time histogram (bin size = 200ms, step size = 2ms, shaded areas represented ± s.e.m. across trials). Black dashed lines indicate stimulus onsets and offsets.

**Extended Data Fig. 9:**
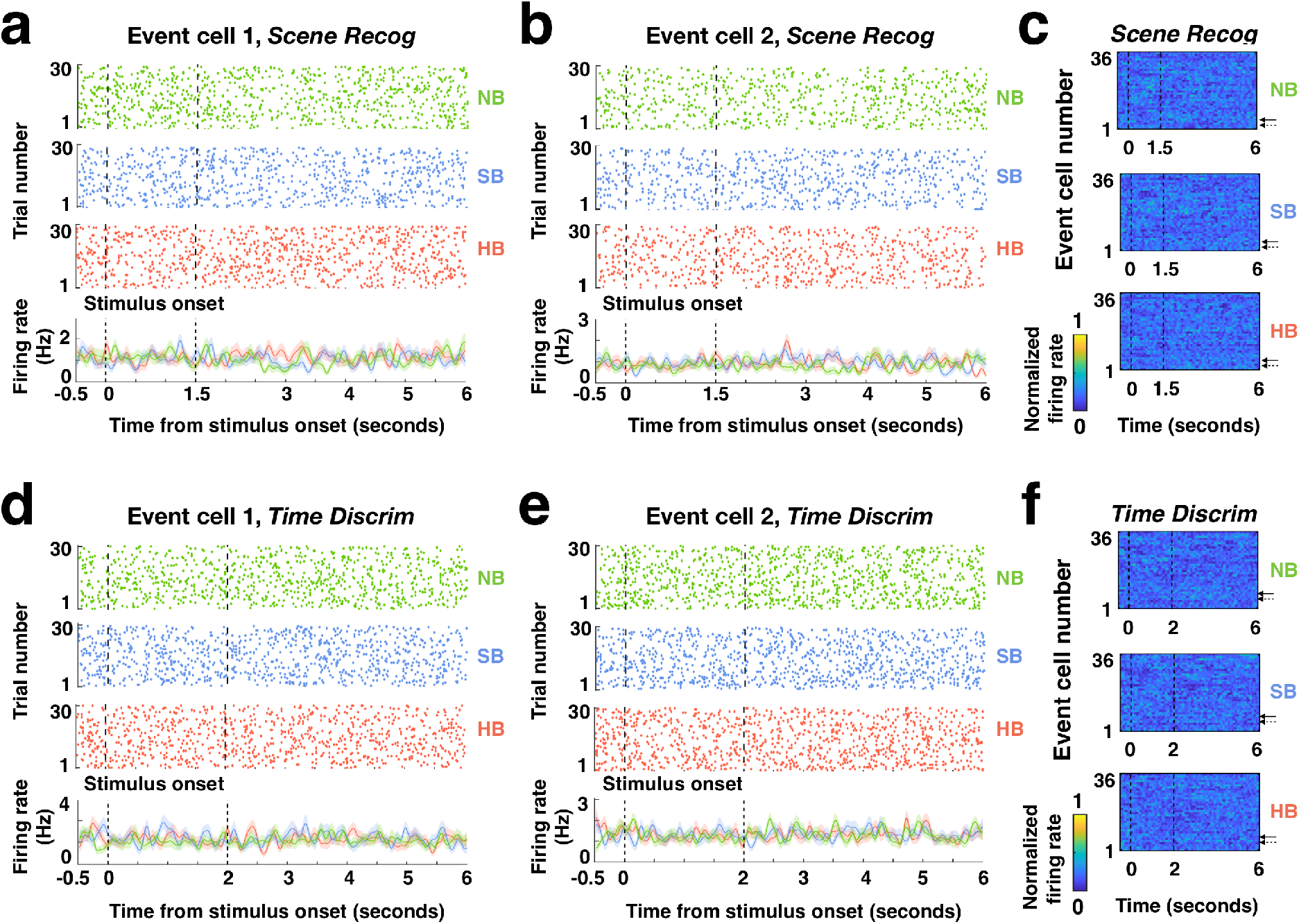
Responses of event cells during scene recognition and time discrimination tasks: **a-b**, Responses during the scene recognition stage from the same example event cells shown in **Fig.3e-f** aligned to the stimulus onsets. **c**, Firing rates of all 36 event cells (solid and dashed arrows denote the examples in **a** and **b**, respectively) during the scene recognition aligned to the stimulus onsets, averaged over trials within each boundary type and normalized to each neuron’s maximum firing rate throughout the entire task (see color scale on bottom). **d-e**, Responses during the time discrimination stage from the same example event cells shown in **Fig.3e-f** aligned to the stimulus onsets. Same format as **a-b**. **f**, Firing rates of all 36 event cells (solid and dashed arrows denote the examples in **d** and **e**, respectively) during the time discrimination stage, using the same format as **c**. For **a, b, d, e**, Top: raster plot color coded for different boundary types (green: NB; blue: SB; red: HB). Bottom: Post-stimulus time histogram (bin size = 200ms, step size = 2ms, shaded areas represented ± s.e.m. across trials). Black dashed lines indicate stimulus onsets and offsets.

**Extended Data Fig. 10:**
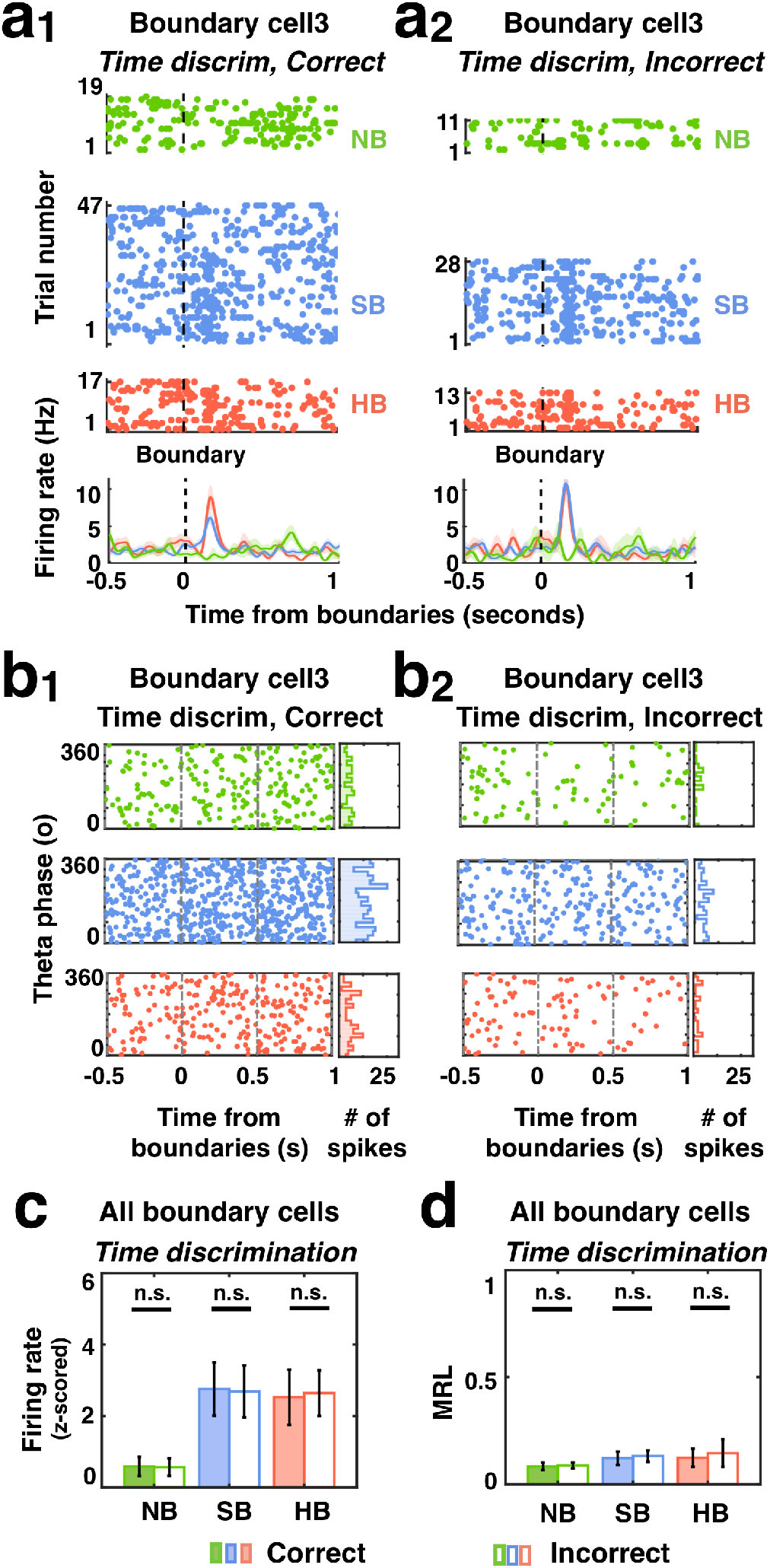
Responses of boundary cells during encoding grouped by memory outcomes from the time discrimination task. **a_1_-a_2_.** Response of the same example boundary cell in **Fig. 4a-b**. During encoding, this cell responded to SB and HB transitions regardless of whether the temporal order of the clip was later correctly (**a_1_**) or incorrectly (**a_2_**) recalled in the time discrimination task. **b_1_-b_2_**. Left: timing of spikes from the same boundary cell shown in **a_1_-a_2_** relative to theta phase calculated from the local field potentials, for clips whose temporal orders were later correctly (**b_1_**) or incorrectly (**b_2_**) recalled. Right: phase distribution of spike times in the 1s period following the middle of the clip (NB) or boundary (SB, HB) for clips whose temporal orders were later correctly (**b_1_**) or incorrectly (**b_2_**) recalled. **c-d.** Population summary for all 42 boundary cells. **c.** Z-scored firing rate (0-1s after boundaries during encoding) for each boundary type did not differ significantly between clips whose temporal orders were later correctly (color filled) vs. incorrectly (empty) recalled. **d**. Mean resultant length (MRL) of spike times (relative to theta phases, 0-1s after boundaries during encoding) across all boundary cells for each boundary type did not differ significantly between clips whose temporal orders were later correctly (color filled) vs. incorrectly (empty) recalled. Error bars indicate standard deviation across n= 42 cells, one-way ANOVA, degrees of freedom = (1, 82).

**Extended Data Fig. 11:**
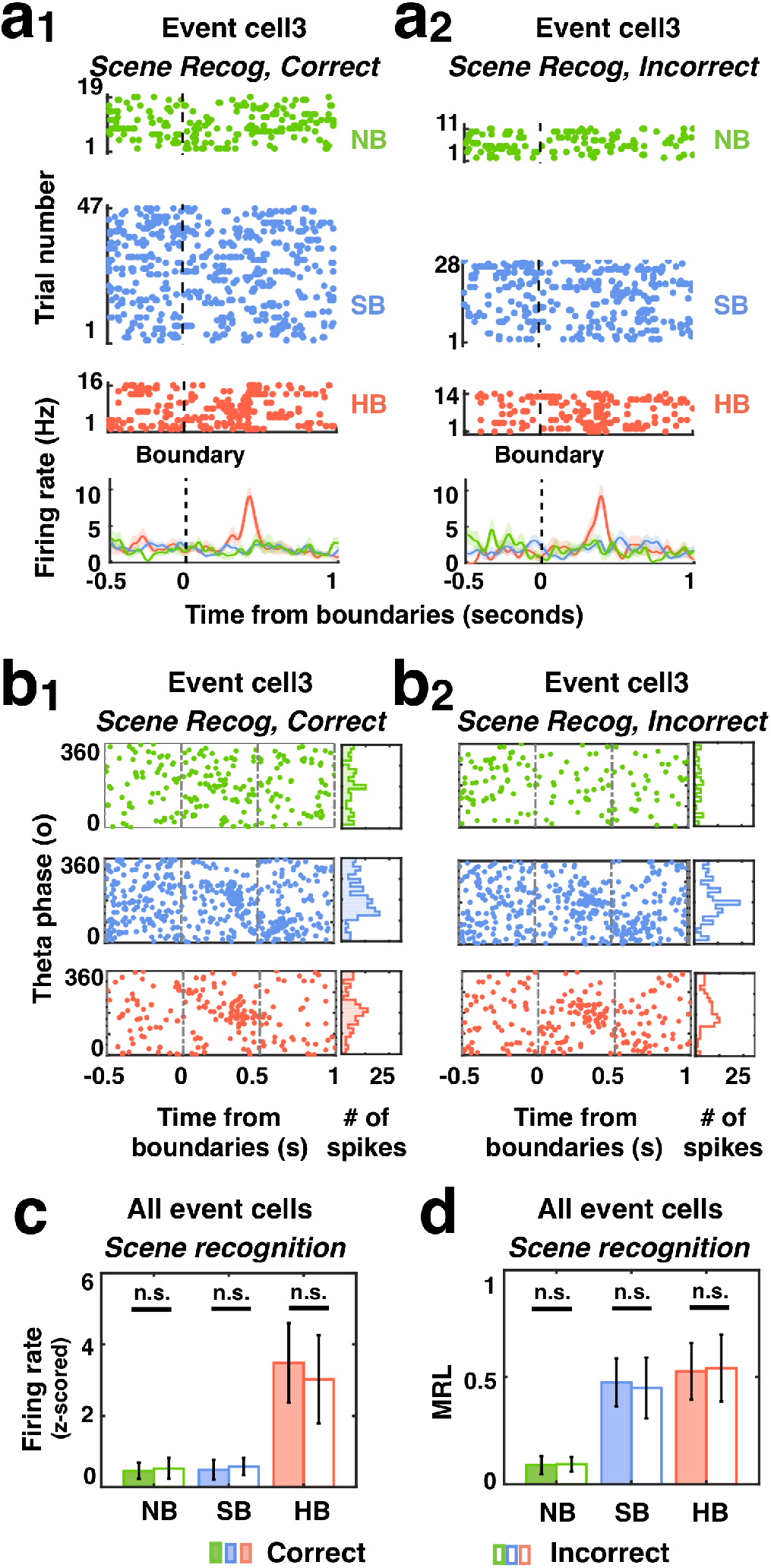
Responses of event cells during encoding grouped by memory outcomes from the scene recognition stage. **a_1_-a_2_.** Response of the same example event cell in **Fig. 4e-f**. During encoding, this cell responded to HB transitions regardless of whether frames were later correctly (**a_1_**) or incorrectly (**a_2_**) recognized in the scene recognition task. **b_1_-b_2_**. Left: timing of spikes from the same event cell shown in **a_1_-a_2_** relative to theta phase calculated from the local field potentials, for frames that were later correctly (**b_1_**) or incorrectly (**b_2_**) recognized. Right: phase distribution of spike times in the 1s period following the middle of the clip (NB) or boundary (SB, HB) for frames that were later correctly (**b_1_**) or incorrectly (**b_2_**) recognized. **c-d.** Population summary for all 36 event cells. **c.** Z-scored firing rate (0-1s after boundaries during encoding) for each boundary type did not differ significantly between frames that were later correctly (color filled) vs. incorrectly (empty) recognized. **d**. Mean resultant length (MRL) of spike times (relative to theta phases, 0-1s after boundaries during encoding) across all event cells for each boundary type did not differ significantly between frames that were later correctly (color filled) vs. incorrectly (empty) recognized. Error bars indicate standard deviation across n =36 cells, one-way ANOVA, degree of freedom = (1, 70).

**Extended Data Fig. 12:**
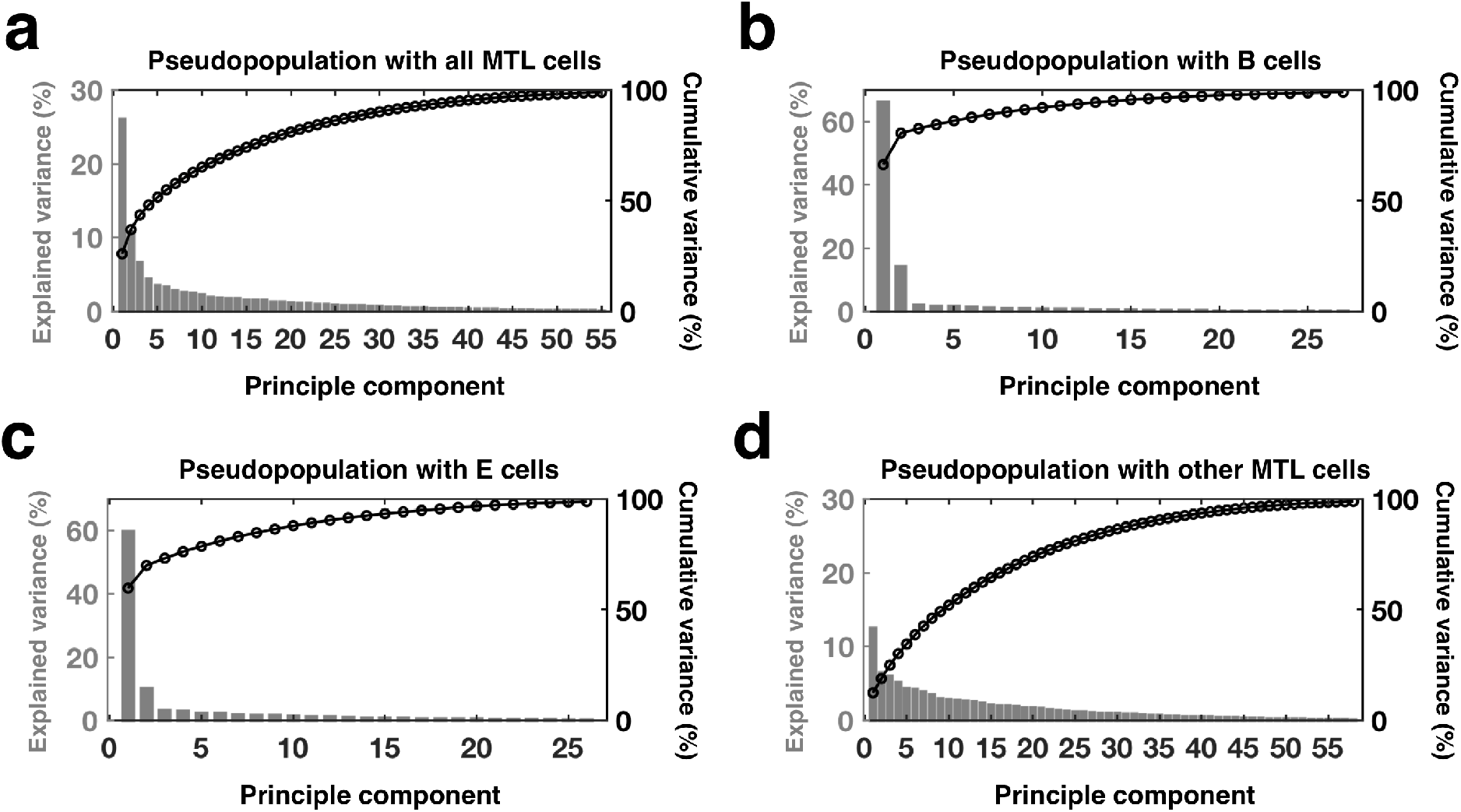
Percentage of explained variance in reconstructed principal component space. Percent variance explained by top principal components (PC) for neural activity from (**a**) all MTL cells, (**b**) boundary cells, (**c**) event cells and (**d**) other MTL cells (i.e., non-boundary/event cells in the MTL). Each bar contains information about the proportion of variance of each and each line denotes the cumulative variances. Full variance (100%) was carried in top 55 PCs in (**a**), top 27 PCs in (**b**) and top 26 PCs in (**c**) and top 58 PCs in (**d**).

**Extended Data Fig. 13:**
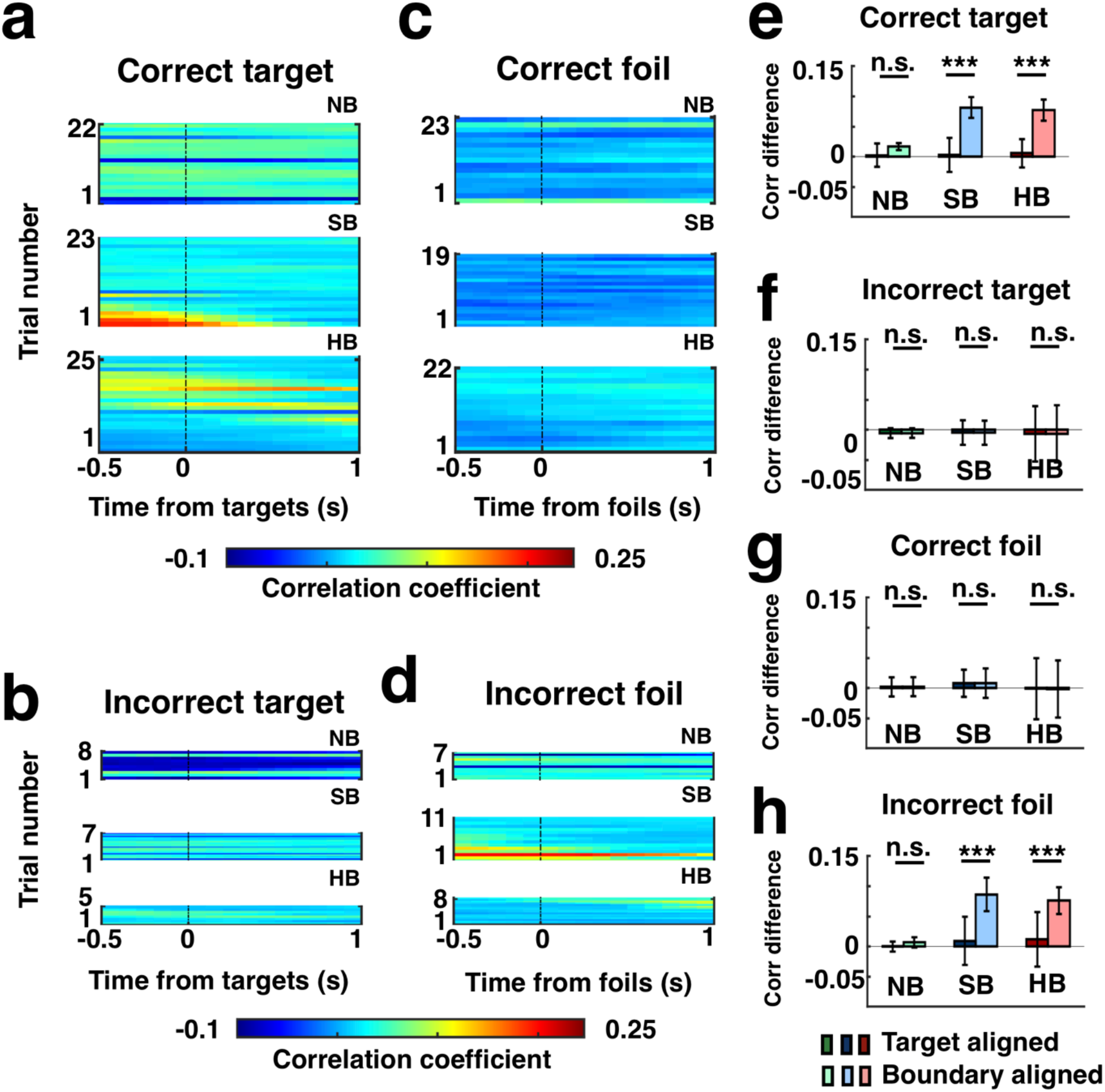
Stronger reinstatement of neural context at boundaries than target: **a-d.** Example subject (same subject as in **Fig 6a-d**) showing trial-by-trial correlation between the population responses during the scene recognition task (0-1.5s relative to stimulus onsets) and during the encoding period (sliding window of 1.5s and 200ms step size), aligned to the target time (time = 0s) for correctly recognized (**a-c**) or incorrectly recognized (**b-d**) target or for foil trials, the time in the clip of their corresponding targets (see Methods). These plots are identical to **Fig. 6a-d**, except here t=0 is where the target was rather than the boundary. **e-h.** Difference of correlation coefficients between [0 1]s and [−1 0]s intervals with respect to targets (dark) or boundaries (light) for each boundary type (green: NB; blue: SB; red: HB), as shown in part **a-d**, then averaged across all the subjects for correctly recognized (**e, g**) or incorrectly recognized (**f, h**) target or foil trials. Error bars indicate standard deviation across n= 19 subjects. \*\*\**P* < 0.001, one-way ANOVA, degrees of freedom = (1, 36).

